# Gibberellin Perception Sensors 1 and 2 reveal cellular GA dynamics articulated by COP1 and GA20ox1 that are necessary but not sufficient to pattern hypocotyl cell elongation

**DOI:** 10.1101/2023.11.06.565859

**Authors:** Jayne Griffiths, Annalisa Rizza, Bijun Tang, Wolf B. Frommer, Alexander M. Jones

## Abstract

The phytohormone gibberellin (GA) is critical for environmentally sensitive plant development including germination, skotomorphogenesis and flowering. The FRET biosensor GIBBERELLIN PERCEPTION SENSOR1, which permits single-cell GA measurements *in vivo*, was previously used to observe a GA gradient correlated with cell length in dark-grown but not light-grown hypocotyls. We sought to understand how light signalling integrates into cellular GA regulation. Here we show how the E3 ligase COP1 and transcription factor HY5 play central roles in directing cellular GA distribution in skoto- and photomorphogenic hypocotyls, respectively. We demonstrate that the expression pattern of biosynthetic enzyme *GA20ox1* is the key determinant of the GA gradient in dark-grown hypocotyls and is a target of COP1 signalling. We engineered a second generation GPS2 biosensor with improved orthogonality and reversibility to show the cellular pattern of GA depletion during the transition to growth in the light. This GA depletion partly explains the resetting of hypocotyl growth dynamics during photomorphogenesis. Achieving cell-level resolution has revealed how GA distributions link environmental conditions with morphology and morphological plasticity and the GPS2 biosensor is an ideal tool for GA studies in further conditions, organs and plant species.

## Introduction

The cellular dynamics of growth regulatory phytohormones act as signal integration nodes that allow plant development to continually adjust to environmental conditions. Gibberellin (GA) hormones are required for cell and organ elongation in many developmental contexts, including several with implications for agricultural yields such as germination, budbreak, stem elongation, branching and flowering (Gao and Chu, 2020; Liu et al., 2022; Cheng et al., 2022; Wu et al., 2021; Wang et al., 2021). Early in the life-cycle of the model plant *Arabidopsis thaliana*, GA mediated organ elongation is highly plastic in response to the light environment. Pre-germination, light triggers GA accumulation that allows elongating radicles to emerge through seed coats, while post-germination, darkness triggers GA accumulation in elongating hypocotyls as part of the skotomorphogenesis program, i.e. etiolation, allowing for seedling emergence from soil. In contrast, post-germination illumination represses GA accumulation and hypocotyl elongation as part of photomorphogenesis or deetiolation (Weller et al., 2009; Zhao et al., 2007; Reid et al., 2002; Alabadí et al., 2008). We developed a genetically-encoded FRET biosensor for GA, nuclear localised Gibberellin Perception Sensor 1 (nlsGPS1), that revealed cellular GA dynamics in skoto- and photomorphogenesis (Rizza et al., 2017). Interestingly, skotomorphogenic hypocotyls showed low GA levels in the apical hook grading to high GA levels in the sub-apical hypocotyl, a cellular dynamic that correlated with cell length patterns. However, it remained unclear how this cellular GA gradient arises, is reprogrammed for photomorphogenesis, and is quantitatively related to cell elongation.

A cellular gradient of a mobile hormone such as GA can emerge from a number of differentially regulated enzymatic steps including early biosynthetic steps leading to the mobile intermediate GA_12_, the penultimate step catalyzed by GIBBERELLIN 20-OXIDASEs (GA20ox), the final biosynthetic step catalyzed by GIBBERELLIN 3-OXIDASEs (GA3ox), and catabolic steps catalyzed by GIBBERELLIN 2-OXIDASES (GA2ox) or CYTOCHROME P450, FAMILY 714, SUBFAMILY A, POLYPEPTIDE 1s (CYP714A1s) (Hedden, 2020; Yamaguchi et al., 2014). In addition to the etiolated hypocotyl gradient, nlsGPS1 also revealed a GA gradient correlated with cell length in the elongation zone of Arabidopsis root tips (Rizza et al., 2017). This finding controverted a computationally simulated root GA gradient based in part on the expression pattern of a GA20ox enzyme as this step was regarded as rate-limiting (Band et al., 2012). Using biosensing, chemical and genetic perturbations, and further mathematical modelling, a key role for the GA3ox step in controlling the elongation zone GA gradient was recently revealed (Rizza et al., 2021). However, it remained unclear which steps are patterned to articulate cellular GA dynamics in the hypocotyl or how they are themselves regulated.

Etiolated hypocotyls express GA20ox1, GA20ox2, GA3ox1, GA2ox1, GA2ox2, GA2ox6, Ga2ox8 and GA2ox9 enzymes, and the expression of several enzymes changes dynamically in response to light during Arabidopsis de-etiolation (Sun et al., 2016). In de-etiolating pea seedlings, red, far-red and blue light inhibit expression of GA3ox genes and lower GA levels within 2-4 hours (Reid et al., 2002). We observed low GA levels without a clear GA gradient in light grown Arabidopsis hypocotyls (Rizza et al., 2017), suggesting that elevated cellular GA levels in the sub-apical hypocotyl would be depleted during de-etiolation. Phytochrome inhibition of hypocotyl elongation is well known to involve lowered GA signalling (Kamiya and García-Martínez, 1999; Hedden and Phillips, 2000; García-Martinez and Gil, 2001; Halliday and Fankhauser, 2003; Vandenbussche et al., 2005). However, a double mutant in Arabidopsis red and far-red-sensing PHYTOCHROME A and PHYTOCHROME B (*phyAphyB*) expressing nlsGPS1 showed only modestly elevated GA levels in the light (Rizza et al., 2017). Blue light-sensing CRYPTOCHROMES (CRY1 and CRY2) are positive regulators of the catabolic *GA2ox* genes and negative regulators of the biosynthesis genes *GA20ox1* and *GA3ox1* in Arabidopsis (Zhao et al., 2007). PHY and CRY signaling both antagonize the central skotomorphogenesis regulator CONSTITUTIVE PHOTOMORPHOGENESIS 1 (COP1), an E3 ubiquitin-ligase with myriad degradation targets (Kim et al., 2017; McNellis et al., 1994). Among these, ELONGATED HYPOCOTYL 5 (HY5) is a transcription factor playing a key role in promoting photomorphogenesis in the light (Xiao et al., 2021; Koornneef et al., 1980). On the other hand, transcription factors promoting skotomorphogenesis include a series of basic helix-loop-helix (bHLH) proteins PHYTOCHROME INTERACTING FACTORS (PIFs) (Leivar and Quail, 2011) and brassinosteroid responsive BRI1-EMS SUPRESSOR1 (BES1) (Yin et al., 2002) and BRASSINAZOLE RESISTANT 1 (BZR1) (Wang et al., 2002). Although PIFs, BES1 and BZR1 are dependent on GA signaling for removal of antagonist DELLA proteins (Feng et al., 2008; de Lucas et al., 2008; Gallego-Bartolomé et al., 2012; Oh et al., 2012; Wang et al., 2012; Bai et al., 2012; Li et al., 2012; Oh et al., 2014), there is also evidence that they can promote GA accumulation in positive feedback loops acting in rapid elongation growth of the sub-apical hypocotyl (Rizza et al., 2017).

Here we investigated the quantitative effects of the above enzymatic steps and light signaling components on the cellular GA dynamics of etiolated hypocotyls. In darkness, PIF and BR signalling influenced GA homeostasis but were not required for establishing the GA gradient while COP1 played a critical role in GA accumulation in the sub-apical hypocotyl. Although GA2ox catabolism contributed to limiting GA from the apical hook, we pinpointed spatial control over GA20ox1 as a key generator of the GA gradient as its expression correlated with cellular GA accumulation in wildtype and mutants and induction of GA20ox1 expression strongly raised GA levels across the etiolated hypocotyl. In the light, PHY and CRY signaling repressed GA accumulation, in part through HY5, and a *phyAphyBcry1cry2* mutant restored the skotomorphogenic GA gradient. In order to observe the effects of deetiolation on patterning of GA, we created a next generation GPS2 biosensor that has improved reversibility over GPS1.

GPS2 was also engineered to have increased orthogonality and showed a loss of GA hypersensitivity phenotypes previously observed in Arabidopsis expressing GPS1. During deetiolation, GA remained low in the apical hook and depleted from the sub-apical hypocotyl, in keeping with lowered biosynthetic and increased catabolic gene expression. Interestingly, a novel *ga2ox* heptuple mutant overelongated during deetiolation. Taken together, our results indicate that GA accumulation is necessary for growth of sub-apical cells but is not sufficient for setting cell length. Additionally, depletion of GA from the apical hook is not developmentally meaningful, but timely GA depletion during deetiolation is important for growth arrest.

## Results

### Skotomorphogenesis agonist PIFs are required to maintain, but not generate GA gradient in dark-grown hypocotyls

We previously demonstrated a reduction of GA levels in a quadruple mutant (*pifq*) lacking four PIFs in the 3 day old dark grown hypocotyl (Rizza et al., 2017) and thus PIFs represented candidate light signalling components that could drive GA accumulation patterns in the etiolated hypocotyl . However, the *pifq* mutant is photomorphogenic with a relatively short hypocotyl and open cotyledons (Leivar et al., 2008) and thus GA regulation could be an indirect consequence of a broader fate switching. To further explore the effect of PIFs on hypocotyl development and GA distribution we monitored the growth of *pifq* and Col-0 in the dark for 72 hours after germination using an infra-red camera. Apical hook development consists of three phases: formation, maintenance and opening (Vandenbussche et al., 2010; Gallego-Bartolomé et al., 2011) .

Surprisingly, we observed the *pifq* mutant completing the formation phase and then opening without entering the maintenance phase (Figure 1a and b). We imaged the upper hypocotyl (0-∼1000 µm from the shoot apical meristem) and analysed GA levels in Col-0 nlsGPS1 and *pifq* nlsGPS1 at 32 hours after germination when Col-0 was still in the maintenance phase and *pifq* was fully open. We observed that *pifq* seedlings have reduced GA levels in the sub-apical hypocotyl and have no apical-basal GA gradient (Figure 1c and e; Figure S1). However, at 24 hours after germination when seedlings were at the end of formation phase, both Col-0 and *pifq* have an apical-basal GA gradient (Figure 1 c and d; Figure S1). The presence of and subsequent loss of the gradient in *pifq* indicates PIFs are not required for the formation of the GA gradient but they are required for its maintenance.

**Figure 1.**
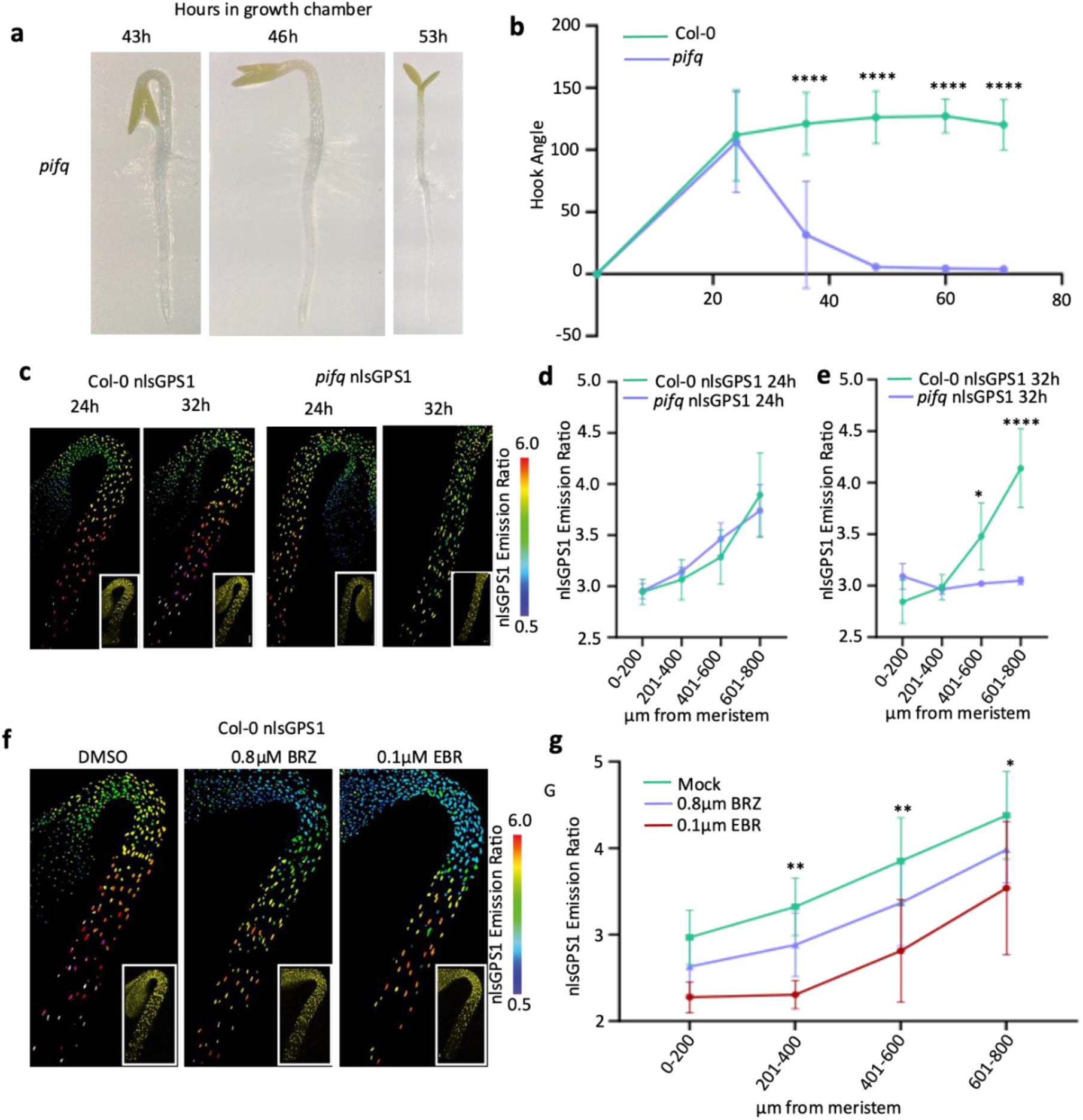
Skotomorphogenesis agonist PIFs are required to maintain, but not generate GA gradient in dark-grown hypocotyls. a) Representative images of *pifq* apical hooks in dark grown seedlings taken at 43h, 46h and 53h after a 4h germination light pulse. b) Apical hook angle in WT Col-0 vs *pifq* mutants. Seedlings grown in the dark were imaged hourly using infrared light in a dark chamber (12h time points shown for simplicity). Repeated measures Two-way ANOVA (Genotype *F* = 142.4, *P* < 0.0001 d.f. = 1; Time *F* = 53.06, *P* < 0.0001 d.f. = 5; Interaction *F* = 35.19, *P* < 0.0001, d.f. = 5; Col-0 *n*=8 *pifq* n=11 biologically independent hypocotyls). A Tukey’s post hoc test was used for multiple comparisons ****p-value < 0.0001. c) nlsGPS1 emission ratios of dark grown hypocotyls 24h and 32h after germination. Representative images of emission ratios and YFP fluorescence (inset) are shown (all images are shown in figure S1). d) Mean of mean nuclear emission ratios from 3 hypocotyls, binning by distance from the shoot apical meristem at 24h after germination. Two-way ANOVA (Genotype *F* = 0.9613, *P* < 0.7605 d.f. = 1; Distance *F* = 17.76, *P* < 0.0001 d.f. = 3; Interaction *F* = 0.5939, *P* =0.6280, d.f. = 3; Col-0 *n* = 3 *pifq* n=3 biologically independent hypocotyls). A Holm-Šídák’s multiple comparisons post hoc test was used for multiple comparisons. e) Mean of mean nuclear emission ratios from 3 hypocotyls, binning by distance from the shoot apical meristem at 32h after germination. Two-way ANOVA (Genotype *F* = 15.88, *P* = 0.0011 d.f. = 1; Distance *F* = 12.71, *P* = 0.0002 d.f. = 3; Interaction *F* = 12.41, *P* = 0.0002, d.f. = 3. (Col-0 *n*=3 *pifq* n=3 biologically independent hypocotyls). A Holm-Šídák’s multiple comparisons post hoc test was used for multiple comparisons *p-value < 0.05, ****p-value < 0.0001. f) nlsGPS1 emission ratios of 3d dark grown hypocotyls treated with 0.1µm epi-brasssonolide (EBR) or 0.8µm brassinazole (BRZ) for 24h before imaging with DMSO treatment as mock. Representative images of emission ratios and YFP fluorescence (inset) are shown (all images are shown in figure S2). g) Quantification of nlsGPS1 emission ratios in response to 24h 0.1µM EBR or 0.8µM BRZ treatment. Mean of mean nuclear emission ratios from 3 hypocotyls, binning by distance from the shoot apical meristem. Two-way ANOVA (Treatment *F* = 16.43, *P* < 0.0001 d.f. = 2; Distance *F* = 21.76, *P* < 0.0001 d.f. = 3; Interaction *F* = 0.1380, *P* = 0.9903, d.f. = 6; *n* = 3 biologically independent hypocotyls). A Tukey’s post hoc test was used for multiple comparisons to mock *p-value < 0.05, **p-value < 0.01.

### Brassinosteroid distribution does not explain GA gradient in dark-grown hypocotyls

Brassinosteroids are known regulators of hypocotyl elongation and GA metabolism making them a candidate for controlling GA patterning. Brassinosteroids have a dual regulatory role on GA levels promoting both GA biosynthesis and GA catabolism (Unterholzner et al., 2015; Albertos et al., 2022) . We have established that the GA gradient is set up early in development. To investigate whether BR could be establishing this gradient, we first reduced BR levels by growing Col-0 nlsGPS1 in the presence of the BR biosynthesis inhibitor brassinazole (BRZ). To avoid germination effects, we germinated Col-0 nlsGPS1 on media and transferred seedlings to BRZ at 48 hours after a germination light pulse. We observed an overall decrease in GA levels, however a clear apical-basal gradient of GA was maintained (Figure 1f and g; Figure S2).

Second, we increased BR levels by growing Col-0 nlsGPS1 in the presence of epibrassinolide (24-epiBL) and again saw an overall decrease of GA levels in the hypocotyl with persistence of the apical-basal gradient (Figure 1f and g; Figure S2). These results support a complex role for brassinosteroids in the regulation of GA levels and indicate that BR levels are optimised in the plant to give high levels of GA that are required for rapid elongation in the dark. However, the persistence of the gradient under altered brassinosteroid levels indicates that brassinosteroids are not responsible for apical-basal patterning of GA.

### Skotomorphogenesis agonist COP1 is required for generating GA gradient and GA20ox1 and GA3ox1 expression in dark-grown hypocotyls

Photoreceptors play key roles in both GA responsiveness and modulating GA levels. CRY1 and PHYA have been shown to redundantly regulate expression of GA metabolic enzymes and to lower GA levels in pea hypocotyls (Reid et al., 2002) and indeed we have previously shown that in Arabidopsis the *phyAphyB* mutant has elevated levels of GA in the light (Rizza et al., 2017). CRYs are positive regulators of catabolic *GA2ox* genes and negative regulators of the biosynthesis genes *GA20ox1* and *GA3ox1* (Zhao et al., 2007). The quadruple *phyAphyBcry1cry2* mutant grown under white light produces a hypocotyl similar in length to dark grown wild-type plants (Mazzella et al., 2001). To further investigate the role of the photoreceptors in establishing the GA gradient we created *phyAphyBcry1cry2* nlsGPS1. Under white light photomorphogenesis is severely delayed in this line and we observed elevated levels of GA forming an apical-basal gradient (Figure 2a and b). As PHYs and CRYs act to reduce GA levels in the light, we explored downstream targets of both CRY and PHY that act to establish the gradient in the dark.

**Figure 2.**
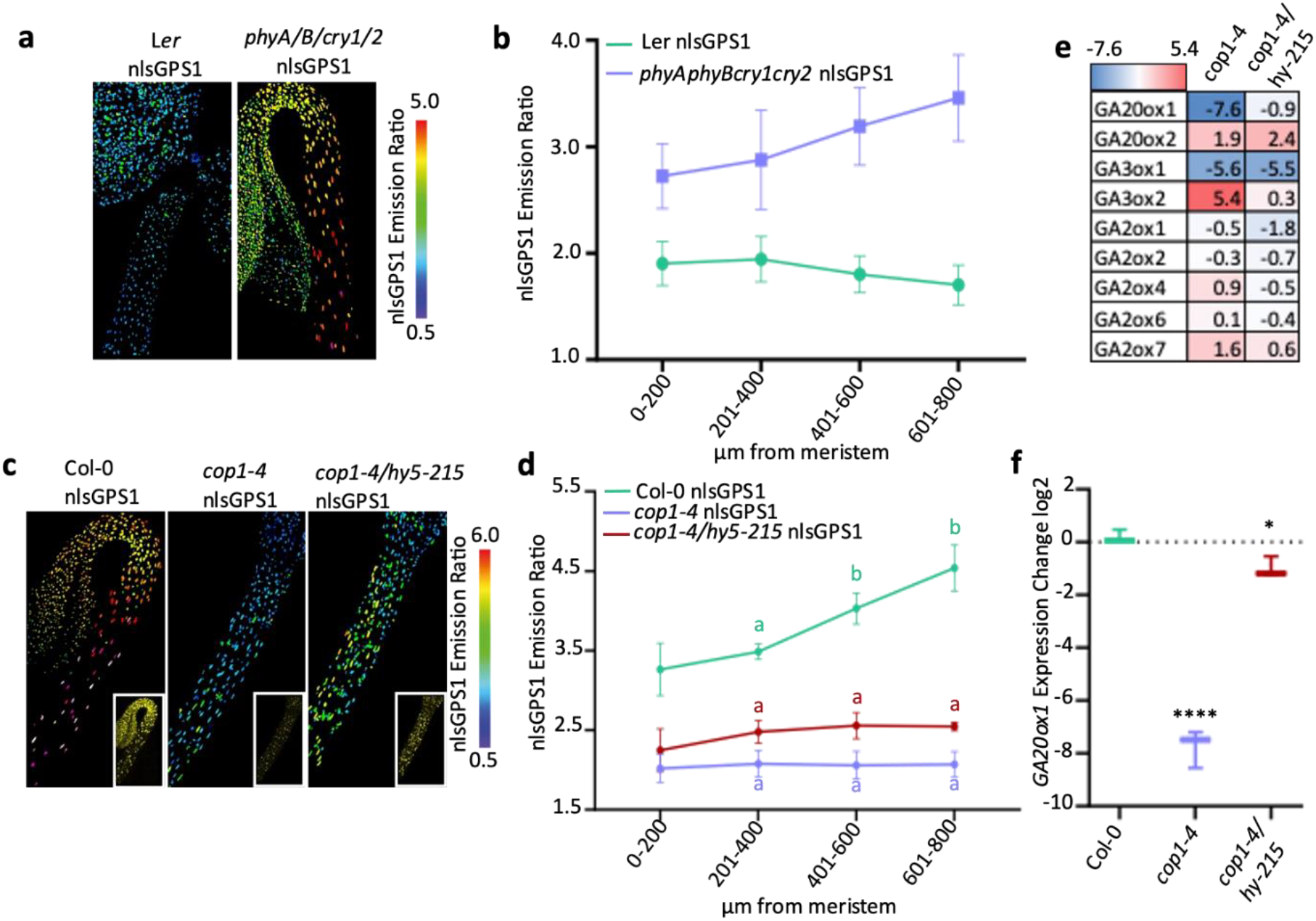
Skotomorphogenesis agonist COP1 is required for GA gradient and *GA20ox1* and *GA3ox1* expression in dark-grown hypocotyls. a) nlsGPS1 emission ratios of 3d white light grown hypocotyls of WT Ler and *phyAphyBcry1cry2* mutant. Representative images of emission ratios are shown. b) Mean of mean nuclear emission ratios from 3 hypocotyls, binning by distance from the shoot apical meristem of 3d dark grown hypocotyls in WT Ler nlsGPS1 and *phyAphyBcry1cry2* nlsGPS1 mutant. c) nlsGPS1 emission ratios of 3d dark grown hypocotyls of WT Col-0 and mutants *cop1-4* and *cop1-4/hy5-215*. Representative images of emission ratios and YFP fluorescence (inset) are shown. d) Mean of mean nuclear emission ratios from 3 hypocotyls, binning by distance from the shoot apical meristem of 3d dark grown hypocotyls for WT Col-0 and mutants *cop1-4* and *cop1-4/hy5-215.* Two-way ANOVA (Genotype *F* = 348.9, *P* < 0.0001 d.f. = 2; Distance *F* = 16.81, *P* < 0.0001 d.f. = 3; Interaction *F* = 9.173, *P* < 0.0001, d.f. = 6. (Col-0 *n*=4, *cop1-4* n=4, *cop1-4/hy5-215* n=4 biologically independent hypocotyls). A Tukey’s multiple comparisons post hoc test was used for multiple comparisons between distance windows comparing to the 0-200µm window, colour of letter corresponds to genotype (a = ns, b= p<0.0001). e) Heat-map showing the fold change of relative expression of GA biosynthetic and catabolic genes in dark grown hypocotyls of *cop1-4* and *cop1-4/hy5* mutants as compared to Col-0. The colors highlight log2 fold expression changes (red: increase; blue: decrease). Values were generated from three biological replicates of 30 hypocotyls measured in three technical repeats (for box and whisker (min max) and statistical significance tests see panel F and figure S4). f) *GA20ox1* expression in dark grown hypocotyls. Log2 fold change from Col-0 for *cop1-4* and *cop1-4/hy5-215*. A one-way ANOVA was performed to compare the effect of genotype on gene expression F= 224.1 P < 0.0001. A Dunnet’s post hoc test was used for multiple comparisons ****p-value < 0.0001 *p-value < 0.05.

The COP1 E3-ligase is inactivated by both photoreceptors and interacts with GA levels and signalling in promoting skotomorphogenesis. As COP1 is required to maintain high levels of GA in dark grown hypocotyls (Weller et al., 2009; Blanco-Touriñán et al., 2020) we explored the possibility that COP1 establishes the spatial distribution of GA. Using *cop1-4* nlsGPS1, we confirmed that GA levels are lower than WT and found that the apical-basal gradient is abrogated in dark-grown seedlings (Figure 2c and d; Figure S3). A primary target of COP1 is the transcription factor HY5, a positive regulator of photomorphogenesis, which accumulates in dark grown *cop1-4* seedlings (Osterlund et al., 2000). As in pea, the *hy5* mutation in the *cop1-4* background partially rescues both the phenotype and the GA levels of *cop1-4* in Arabidopsis (Figure 2c; Figure S3) (Weller et al., 2009). However, using nlsGPS1 we observed that this rescue does not extend to the patterning of GA (Figure 2d). Together these results indicate that COP1 functions to prevent HY5 from repressing GA accumulation, though this function alone does not explain the central role for COP1 in regulating the GA gradient.

GA biosynthetic enzymes are expressed in a tissue specific manner (Plackett et al., 2012; Mitchum et al., 2006) and the complexity of GA metabolism provides many candidate regulators of the hypocotyl GA gradient. We previously showed the key regulatory steps controlling a GA gradient in light grown roots vary along the apical-basal axis (Rizza et al., 2021). In order to identify the specific GA metabolism targets of the COP1/HY5 pathway, we carried out qPCR on 3 day old dark grown hypocotyls of WT, *cop1-4* and *cop1-4/hy5-215* (Figure 2e; Figure S4). We found that the genes involved in biosynthesis displayed greater changes than those involved in catabolism with *GA3ox1* and *GA20ox1* particularly downregulated in *cop1-4* dark grown hypocotyls (Figure 2e; Figure S4). Interestingly, only the expression of *GA20ox1* is partially rescued in the *cop1-4/hy5-215* mutant, suggesting that *GA3ox1* is not a primary target of HY5 repression (Figure 2f; Figure S4).

### GA20ox1 and GA3ox1 expression are correlated with GA gradient

To determine if differential expression of *GA20ox1* and *GA3ox1* are involved in establishing the GA gradient we mapped the expression pattern of these two enzymes using pGA3ox1-TC-GUS and pGA20ox1-TL-GUS reporters (Mitchum et al., 2006; Plackett et al., 2012). In etiolated hypocotyls, expression of the two biosynthetic genes is high in the sub-apical hypocotyl and limited in the apical hook (Figure 3a). To validate the expression observed using the reporter lines, qPCR was carried out in dissected regions defined as the apical hook 0-400µm and the sub-apical hypocotyl 400-800µm (Figure 3b). The transcript levels of the two genes mirrors the GUS staining with higher expression present in the sub-apical hypocotyl relative to the apical hook (Figure 3c), and thus both a potentially responsible for generating the GA gradient.

**Figure 3.**
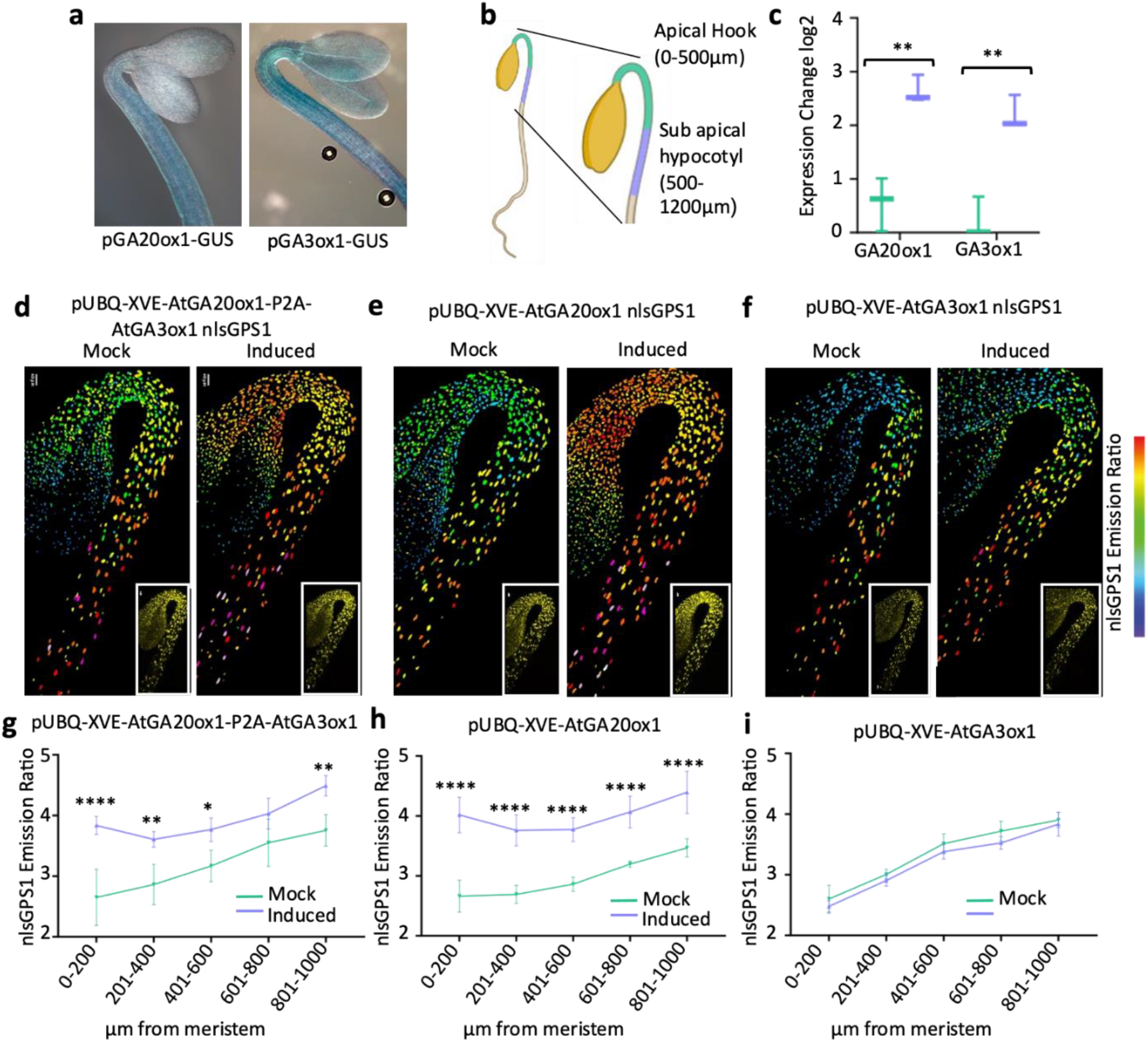
GA3ox1 and GA20ox1 expression are correlated with GA gradient but only GA20ox1 expression is rate limiting in dark-grown hypocotyls. a) 3d dark grown hypocotyls expressing pGA20ox1-GUS translational fusion and pGA3ox1-GUS is a transcriptional fusion from (Plackett et al, 2012 and Mitchum et al., 2008, respectively). b) Schematic diagram showing the position of the apical hook and the sub-apical hypocotyl dissected for qPCR analysis, adapted from “Seedling 1”, by BioRender.com (2023). Retrieved from https://app.biorender.com/biorender-templates. c) Relative expression changes of *GA3ox1* and *GA20ox1* in the apical hook and sub apical hypocotyl. Plants were grown for 3 d in the dark and gene expression analyzed by RT-qPCR. The means and standard deviations of three biological duplicates measured in three technical replicates are plotted. Statistical significance was determined with a Student’s *t* test \*\**P* < 0.01. d-i) nlsGPS1 emission ratios of 3d dark grown hypocotyls β-estradiol inducible GA enzyme transgenic lines 48 h after induction with 2.5 µM 17-β-estradiol (induced) or with 0.1% DMSO mock induction (mock). d-f) Representative images of nlsGPS1 emission ratios and YFP fluorescence (inset) are shown (all images are shown in figure S5). g-i) Mean of mean nuclear emission ratios from 4 hypocotyls, binning by distance from the shoot apical meristem. f) Two-way ANOVA Treatment *F* = 71.95, *P* < 0.0001 d.f. = 1; Distance *F* = 14.95, *P* < 0.0001 d.f. = 4; Interaction *F* = 1.823, *P* = 0.1504, d.f. = 4. g) Two-way ANOVA Treatment *F* = 201.9, *P* < 0.0001 d.f. = 1; Distance *F* = 12.91, *P* < 0.0001 d.f. = 4; Interaction *F* = 1.514, *P* = 0.2233, d.f. = 4. h) Two-way ANOVA Treatment *F* = 7.341, *P* < 0.0110 d.f. = 1; Distance *F* = 110.2, *P* < 0.0001 d.f. = 4; Interaction *F* = 0.2052, *P* = 0.9335, d.f. = 4. Mock n=4, Induced n=4 biologically independent hypocotyls. A Šídák’s multiple comparisons post hoc test was used for multiple comparisons between mock and induced ****p-value < 0.0001, **p-value < 0.01, *p-value < 0.05. Experiments were completed three times with similar results.

### GA20ox1 is a key regulator of GA levels in the dark

In roots, GA20ox and GA3ox enzymatic steps have locally different roles with GA3ox being rate limiting in the elongation zone and both GA20ox and GA3ox being rate limiting in the root initials in the meristem (Rizza et al., 2021). To determine whether the expression of *GA20ox1* and *GA3ox1* may explain the GA gradient in the hypocotyl, we analysed the effect of simultaneous β-estradiol induction of ubiquitous *AtGA20ox1* and *AtGA3ox1* and observed an increase of emission ratio along the hypocotyl with the greatest increase occurring in the apical hook (Figure 3d and g; Figure S5). Lines that express *GA20ox1* or *GA3ox1* individually were used to further dissect the relationship between these GA biosynthetic enzymes and the GA gradient. Induction of *GA20ox1* increased the emission ratio of nlsGPS1 in a similar manner to that observed during simultaneous induction whereas induction of *GA3ox1* alone did not (Figure 3e - i; Figure S5). As an increase of *GA20ox1* expression was sufficient to increase GA levels, *GA20ox1* is alone rate-limiting for GA accumulation in the dark grown hypocotyl, indicating that the expression pattern of this enzyme is a primary driver of the GA gradient here.

### Disruption of the GA gradient is not sufficient to abolish cell length patterning

The *ga20ox1/2/3* mutant has severe developmental defects associated with low levels of GA (Plackett et al., 2012) and we observed that hypocotyl elongation in the dark was dramatically reduced (Figure 4a) as were the levels of GA (Figure 4b; Figure S6). To explore the effect of spatial distribution of GA on cell size we measured epidermal cell lengths in Col-0 nlsGPS1, *ga20ox1/2/3* nlsGPS1 and induced pUBQ-XVE-AtGA20ox1 lines. Cells in the apical hook are smaller than cells in the sub-apical hypocotyl (Gendreau et al., 1997) and this correlates with GA levels (Rizza et al., 2017). As expected, we observed Col-0 cell lengths increasing with distance from meristem in hypocotyls of dark-grown seedlings (Figure 4c). Interestingly, cell length remains patterned when the GA gradient is removed in *ga20ox1/2/3* or when GA levels are raised by inducing ubiquitous expression of *GA20ox1* (Figure 4c). Sub-apical hypocotyl cells are shorter in *ga20ox1/2/3* indicating the level of GA in these cells is important for elongation.

**Figure 4.**
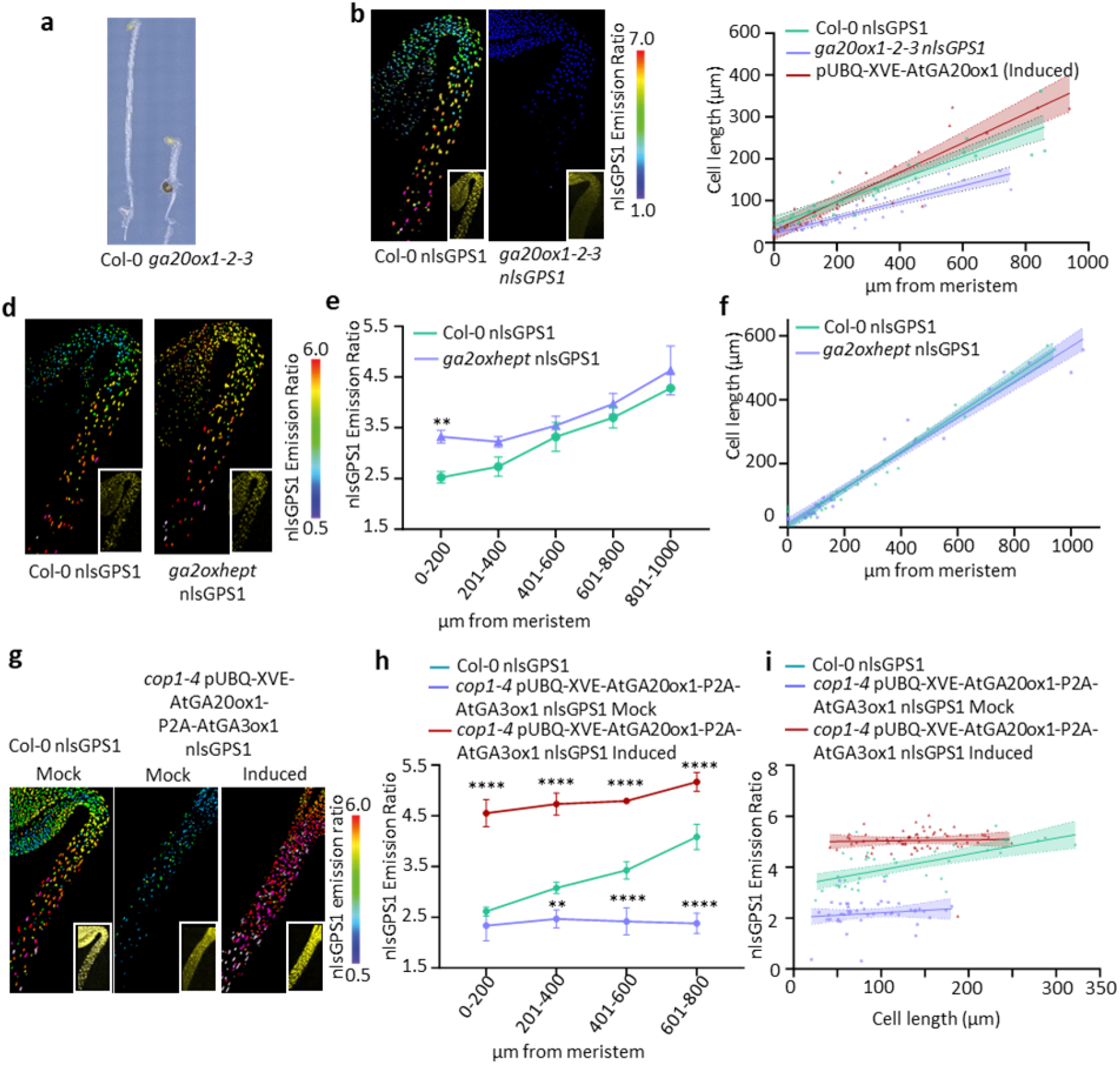
GA catabolism also plays a role in generating the GA gradient in dark-grown hypocotyls, but abolishing the GA gradient is not sufficient to abolish cell length patterning. a) Image of 3d dark grown *ga20ox1,2,3* triple mutant and Col-0 hypocotyls. b) Representative images of nlsGPS1emission ratios and YFP fluorescence (inset) 3d dark grown hypocotyls for WT Col-0 nlsGPS1 and *ga20ox1,2,3* nlsGPS1 are shown (all images are shown in figure S6). c) Scatter plot showing cell length and distance from meristem for individual hypocotyl epidermal cells from the shoot apical meristem. Simple linear regression with 95% confidence intervals plotted. WT Col-0, *ga20ox1,2,3* triple mutant and Col-0 pUBQ-XVE-GA20ox1 line induced with 2.5 µM 17-β-estradiol for 48 h (induced). For Col-0 hypocotyls, Pearson’s correlation coefficient r = 0.92; n=26 cells from 3 hypocotyls (1 cell file per hypocotyl); ga20ox1,2,3 r=0.91, n=35 cells from 3 hypocotyls; pUBQ-XVE-GA20ox1 (induced) r=0.92 n= 27 cells from 3 hypocotyls. d) nlsGPS1 emission ratios of 3d dark grown hypocotyls for WT Col-0 and *ga2oxhept* mutant. Representative images of emission ratios and YFP fluorescence (inset) are shown (all images are shown in figure S7). e) Mean of mean nuclear emission ratios from 3 hypocotyls, binning by distance from the shoot apical meristem of 3d dark grown hypocotyls of WT Col-0 and mutant *ga2oxhept*. Two-way ANOVA (Genotype F = 26.86, P <0.001 d.f. = 1; Distance F = 48.20, P < 0.0001 d.f. = 4; Interaction F = 1.595, P = 0.2146, d.f. = 4. (Col-0 n=3, *ga2oxhept* n=3 biologically independent hypocotyls). A Šídák’s multiple comparisons post hoc test was used for multiple comparisons **p-value < 0.01. f) Scatter plot showing cell length and distance from meristem for individual hypocotyl epidermal cells from the shoot apical meristem. Simple linear regression with 95% confidence intervals plotted. For Col-0 hypocotyls, Pearson’s correlation coefficient r = 0.9849; n=31 cells from 3 hypocotyls (1 cell file per hypocotyl); *ga2oxhept* r=0.9808, n=29 cells from 3 hypocotyls. Experiment was repeated three independent times with similar results. g) Representative images of emission ratios and YFP fluorescence (inset) are shown nlsGPS1 emission ratios of 3d dark grown hypocotyls for WT Col-0 nlsGPS1 and *cop1-4* pUBQ-XVE-AtGA20ox1-P2A-AtGA3ox1 nlsGPS1 (all images are shown in figure S9). β-estradiol inducible GA enzyme transgenic lines 48 h after induction with 2.5 µM 17-β-estradiol (induced) or with 0.1% DMSO mock induction (mock). h) Mean of mean nuclear emission ratios from 3 hypocotyls, binning by distance from the shoot apical meristem of 3d dark grown hypocotyls of WT Col-0, β-estradiol inducible *cop1-4* pUBQ-XVE-AtGA20ox1-P2A-AtGA3ox1 nlsGPS1 mock and induced. Two-way ANOVA (Genotype F = 429.7, P < 0.0001 d.f. = 2; Distance F = 18.89, P < 0.0001 d.f. = 3; Interaction F = 6.999, P = 0.0002, d.f. = 6. (Col-0 n=3, *cop1-4* pUBQ-XVE-AtGA20ox1-P2A-AtGA3ox1 nlsGPS1 mock n=3 induced n=3 biologically independent hypocotyls). A Dunnett’s multiple comparisons post hoc test was used for multiple comparisons comparing to Col-0 control. **p-value < 0.01 ****p-value <0.0001. i) Scatter plot showing cell length and distance from meristem for individual hypocotyl epidermal cells from the shoot apical meristem. Simple linear regression with 95% confidence intervals plotted. For Col-0 nlsGPS1 hypocotyls, Pearson’s correlation coefficient r = 0.65; n=36 cells from 3 hypocotyls. For *cop1-4* pUBQ-XVE-AtGA20ox1-P2A-AtGA3ox1 nlsGPS1 correlation coefficient r = 0.14 and 0.04; n= 46 and 65 cells for mock and induced, respectively.

However, higher levels of GA in the apical hook region do not correlate to longer cells suggesting the relative depletion of GA is not important to keep these cells small. In light grown hypocotyls, where higher GA levels do stimulate cell elongation, GA response was also shown to be positionally dependent along the apical-basal axis (Robinson et al., 2017). Our results indicate that the correlation between GA levels and cell length, as in the root (Rizza et al., 2021), is not quantitatively causative in a simple dose-response relationship.

### GA catabolism also plays a role in generating the GA gradient in dark-grown hypocotyls

Nonetheless, it is intriguing that with the ubiquitous elevation of rate limiting *GA20ox1* expression, GA is slightly lower in the apical hook region compared with the sub-apical hypocotyl. The catabolic GA2ox family consists of seven canonical members and two non-canonical members (Rieu et al., 2008; Schomburg et al., 2003; Lange et al., 2020). Expression data indicates that canonical *GA2ox1,2,4,6,7* and *8* are expressed in etiolating or deetiolating hypocotyls (Sun et al., 2016). In order to determine whether these enzymes are in part responsible for the lowering of GA in the hook we created a *ga2oxheptuple* mutant (*ga2ox1/2/3/4/6/7/8*). Analysis of nlsGPS1 emission ratios in this line indicated increased GA levels in comparison to Col-0 with more GA being present in the apical hook (Figure 4d and e). A lesser apical-basal gradient remains, likely because of patterned expression of *GA20ox1*. As with induced *GA20ox1*, the increased level of GA does not affect the pattern of cell elongation of smaller cells nearer the meristem (Figure 4f), confirming that the correlation of GA levels to cell length observed in the dark grown hypocotyl is not alone causative for growth patterning.

### Biosynthesis is key for GA patterning in *cop1* mutant

We have demonstrated that GA biosynthesis plays a key role in the formation of the apical-basal gradient with *GA20ox1* and *GA3ox1* being key targets of the COP1 pathway. In order to confirm that the short hypocotyl observed in *cop1-4* is due to low GA caused by an impairment of the biosynthetic pathway we created *cop1-4* pUBQ-XVE-AtGA20ox1-P2A-AtGA3ox1 to permit simultaneous β-estradiol induction of ubiquitous *AtGA20ox1* and *AtGA3ox1* in the *cop1-4* background. Hypocotyl length increased with simultaneous β-estradiol induction of ubiquitous *GA20ox1* and *GA3ox1* expression to a similar level as *cop1-4* grown on 1 µM GA_4_ (Figure S8a and b). The increased GA levels and minimal gradient observed in induced *cop1-4* pUBQ-XVE-AtGA20ox1-P2A-AtGA3ox1 confirms that COP1, likely indirectly, influencing *GA20ox1* and *GA3ox1* expression patterns directs cellular GA distribution (Figure 4g and h; Figure S9).

Interestingly, in the *cop1-4* background, both mock and β-estradiol induced pUBQ-XVE-AtGA20ox1-P2A-AtGA3ox1 hypocotyl cells exhibited cell length patterning without correlation with cellular GA levels (Figure 4i). Nonetheless, overall cell lengths were longer in the latter (Figure 4i).

### Next-generation Gibberellin Perception Sensor 2 is reversible and orthogonalized

Although GPS1 has high affinity to bioactive GA and reveals cellular distribution and regulation of GA, slow reversibility precluded use for investigation of GA depletion dynamics (Rizza et al., 2017). Both properties are potentially related to the mechanism of GPS1 biosensing involving GA binding the AtGID1C moiety which then interacts with the truncated DELLA moiety from AtGAI (Figure 5a). Also, the stable transgenic lines expressing nlsGPS1 were hyposensitive to GA biosynthesis inhibitor Paclobutrazol (Rizza et al., 2017), indicating interaction between GPS1 and endogenous GA signaling, likely the AtGID1C moiety with endogenous DELLA proteins (Figure 5a). As FRET biosensor re-engineering can deliver improved variants (e.g. (Rowe et al., 2023)), we aimed to increase GPS1 reversibility and orthogonality, i.e. reduce potential interference with and by endogenous signalling, via re-engineering the GID1-DELLA interaction interface. The high-resolution structure of the *At*GID1A-*At*GAI complex (Shimada et al., 2008) revealed two potential interfacial electrostatic interactions that were conserved in GPS1 as Glutamate 13 (E13) on *At*GID1C and Lysine 51 (K51) on *At*GAI and Glutamate 28 (E28) on *At*GID1C and Arginine 54 on *At*GAI (R54) (Figure 5b). Introducing charge exchange mutations in the AtGAI moiety (K51E and R54E), which would disrupt the inter-domain interaction, was successfully used to engineer the GPS1-NR negative control sensor (Rizza et al., 2017). We introduced four charge exchange mutations E13K and E28K (*At*GID1C) and K51E and R54E (*At*GAI) into GPS1 to restore the original two electrostatic interactions while abrogating interactions between GPS1 and endogenous GID1s and DELLAs (Figure 5c and d).

**Figure 5.**
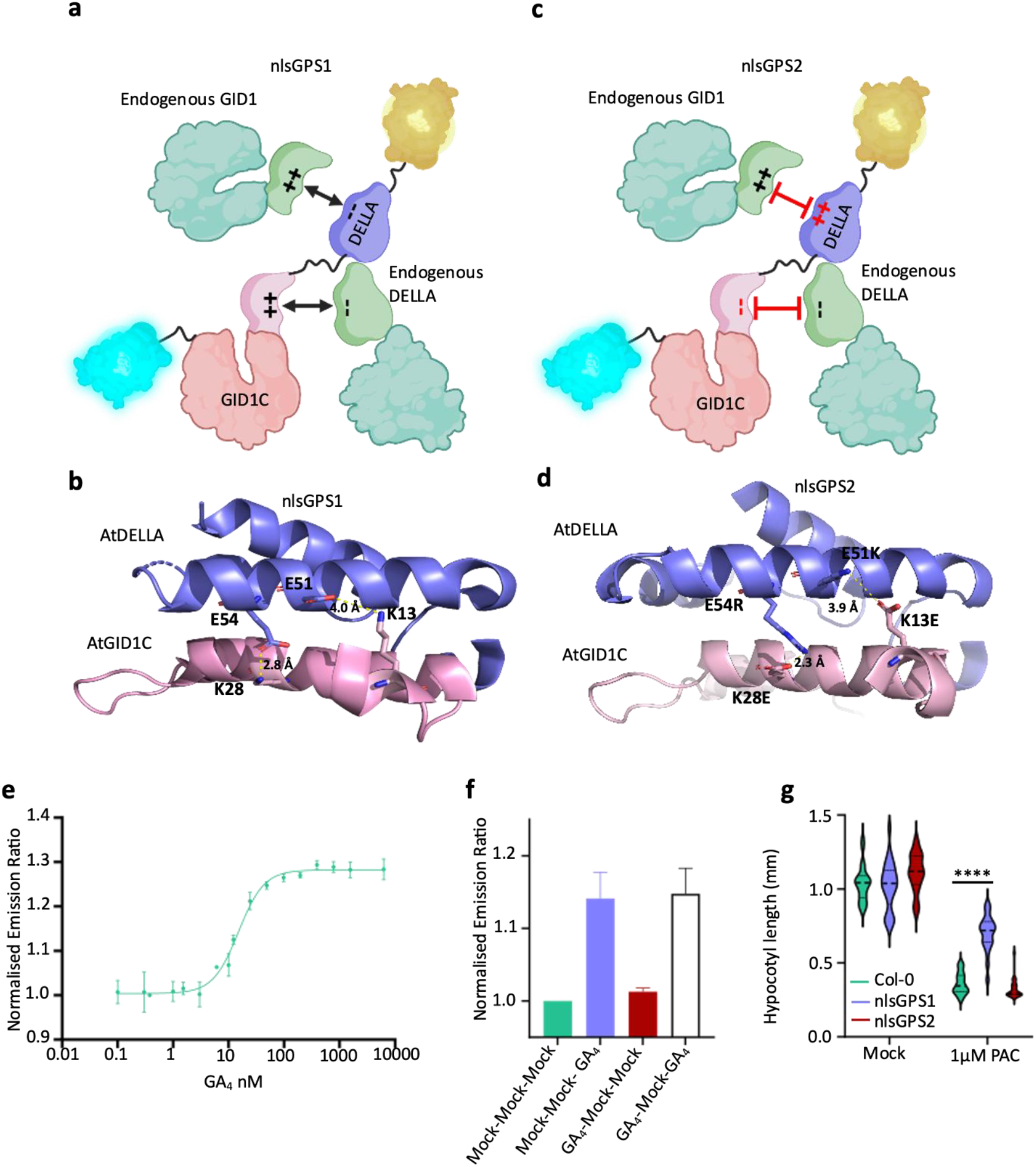
Engineering the orthogonalized and more reversible GPS2 biosensor. a) Graphic of interaction of charged amino acids within the sensory domain of GPS1 with endogenous components (BioRender.com). b) AlphaFold-predicted structure of the GPS1 interface between GAI and GID1C illustrates the presence of charged residues, E51 and E54 (GAI), and K13 and K28 (GID1C), as well as the presence of two predicted electrostatic interactions. c) Graphic of the design strategy for GPS2, involving the exchange of two pairs of charged residues between GID1C and DELLA in the sensory domain (BioRender.com). d) AlphaFold-predicted structure of the GPS2 maintains the electrostatic interactions between the charged residues after undergoing charge-exchange mutations (GAI E51K and E54R, GID1C K13E and K28E). e) Titration of GPS2 purified protein in response to GA_4_. Changes in the ratio were calculated for GPS2 purified protein with varying GA_4_ concentrations. Error bars represent the variation observed across five biological replicates. The dissociation constant (Kd) was calculated as the mean of all biological replicates (n=5). f) *In vitro* reversibility on purified GPS2 protein. Ratio changes were recorded for two groups, one with GA pre-treatment (GA_4_) and one without (Mock). In both groups, the protein was washed through a desalting column twice before GA_4_ treatment (n=2). g) Violin plot of hypocotyl lengths of nlsGPS1 and nlsGPS2 lines were compared with wild-type Col-0 with and without Paclobutrazol (1 µM PAC) treatment. Student’s *t*-test Col-0 to nlsGPS1 and nlsGPS2, \*\*\*\**P* value < 0.0001.

*In vitro* analysis of GPS1 purified from yeast revealed successful restoration of GA emission ratio response and maintenance of a high affinity for bioactive GA_4_ (Figure 5d). *In vitro* reversibility testing using buffer-exchange chromatography revealed that this quadruple charge exchange mutant detected a second GA treatment following a first (Figure 5e). Thus, this biosensor had improved reversibility, likely due to increased off-rate of the GID1-DELLA interaction resulting from inversion of the interfacial hydrostatic interactions. Transgenic Arabidopsis lines constitutively expressing a nuclear localized variant of this biosensor were found to have lost the PAC hyposensitivity phenotypes of nlsGPS1 (Figure 5f). We thus consider the variant to be second generation Gibberellin Perception Sensor 2 (GPS2) with improved reversibility and orthogonality.

### Depletion of hypocotyl GA is detected with nlsGPS2 biosensors

Previous studies have demonstrated that when a hypocotyl underdoes de-etiolation there is an increased expression of GA catabolism genes and a decreased expression of some GA biosynthesis genes which correlates to a drop in bioactive GA (Reid et al., 2002; Zhao et al., 2007; Weller et al., 2009). However, there is also a transient increase of some GA biosynthesis genes which could indicate localised GA increase e.g. in the apical hook. How these expression dynamics affect GA dynamics and whether this is relevant to the deetiolation response was not known. To determine whether the next generation biosensor could be used to image this decrease of GA, we examined how nlsGPS1 and nlsGPS2 respond during de-etiolation. We imaged the non-reversible Col-0 nlsGPS1 and the reversible Col-0 nlsGPS2 6 hours after transfer to the light in comparison with hypocotyls that had been kept in the dark. Col-0 nlsGPS1 did not show a reduction in GA levels, which is contrary to published studies but expected due to the low reversibility the biosensor (Figure 6a and b; Figure S10). One the other hand, in Col-0 nlsGPS2 we observed a reduction in emission ratios along the length of the hypocotyl with the greatest reduction occurring in cells that had high GA levels in the dark (>600µm from the meristem) (Figure 6a and b; Figure S10). Neither sensor reported any localised GA increases at this time point.

**Figure 6.**
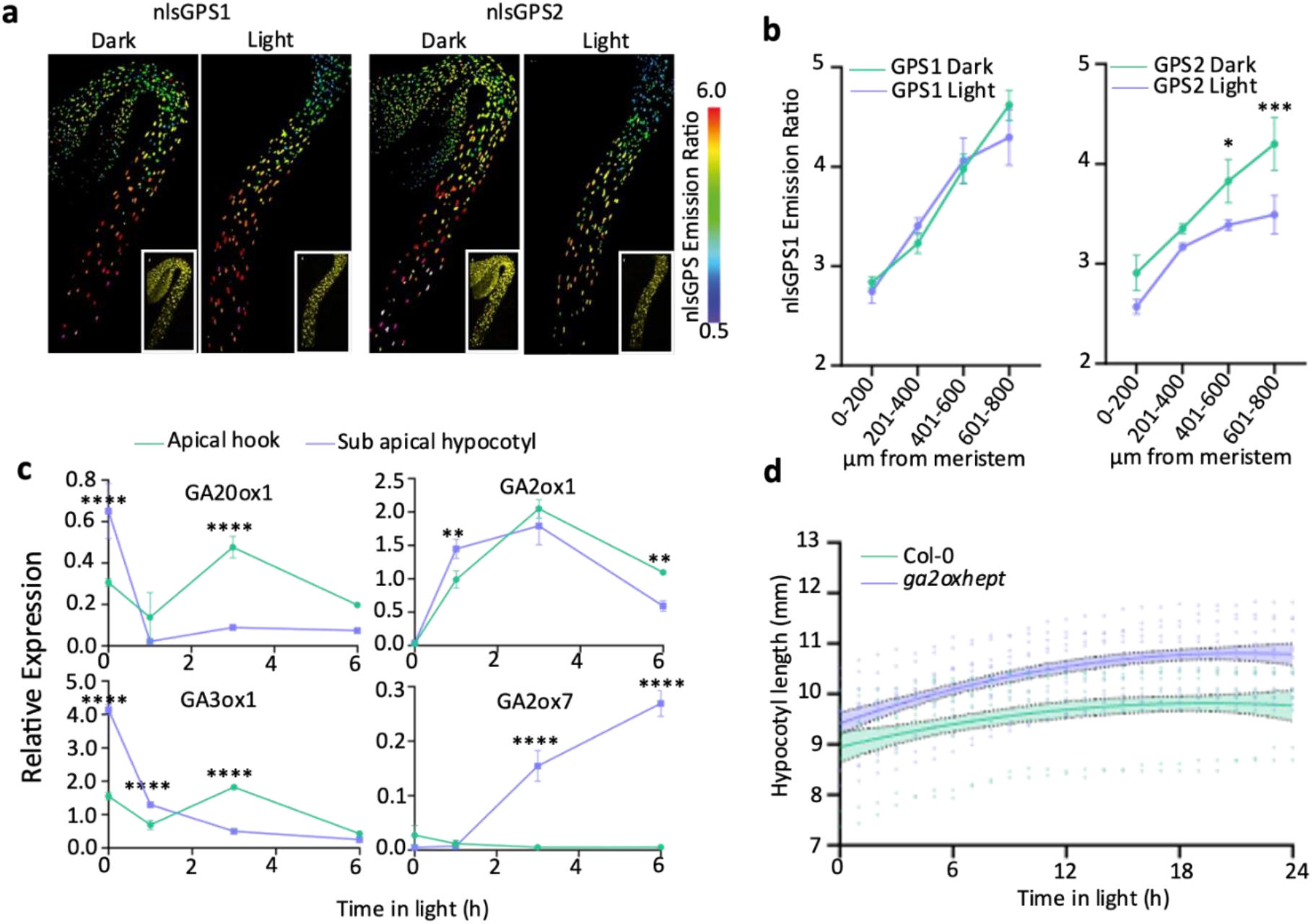
Depletion of hypocotyl GA detectable with nlsGPS2 biosensors and limited GA depletion in a *ga2oxhept* mutant causes hypocotyl hyperelongation during photomorphogenesis. a) Emission ratios of nlsGPS1 and nlsGPS2 in 3d dark grown hypocotyls transferred to light or maintained in darkness for 6h. Representative images of emission ratios and YFP fluorescence (inset) are shown (all images are shown in figure S10). b) Mean of mean nuclear emission ratios from 3 hypocotyls, binning by distance from the shoot apical meristem of 3d dark grown hypocotyls of nlsGPS1 and nlsGPS2 transferred to light for 6 hours or maintained in darkness. nlsGPS1 Two-way ANOVA (Light condition *F* = 0.3392, *P* = 0.5684 d.f. = 1; Distance *F* = 126.0, *P* < 0.0001 d.f. = 3; Interaction *F* = 2.805, *P* = 0.0731, d.f. = 3. A Šídák’s multiple comparisons post hoc test was used for multiple comparisons. nlsGPS2 Two-way ANOVA (Light condition *F* = 42.97, *P* < 0.0001 d.f. = 1; Distance *F* = 56.90, *P* < 0.0001 d.f. = 3; Interaction *F* = 2.805, *P* = 0.0629, d.f. = 3. A Šídák’s multiple comparisons post hoc test was used for multiple comparisons *p-value < 0.05, ***p-value < 0.001. c) Relative expression changes of selected GA metabolic enzymes in the hook and sub-apical hook of Col-0 during deetiolation. Dissections were carried out in the dark and at 1h, 3h and 6h light exposure. Data represents mean and standard deviation of three technical repeats from one biological repeat containing (30) dissected hypocotyls per sample. Experiment was carried out two independent times with similar results (Second replicate shown in figure S11). Two-way ANOVA: GA20ox1 (Tissue *F* = 0.6.715, *P* = 0.0197 d.f. = 1; Time *F* = 43.21, *P* < 0.0001 d.f. = 3; Interaction *F* =31.54, *P* < 0.0001, d.f. = 3). GA3ox1 (Tissue *F* = 131.4, *P* < 0.0001, d.f. = 1; Time *F* = 835.3, *P* < 0.0001 d.f. = 3; Interaction *F* =495.6, *P* < 0.0001, d.f. = 3). GA2ox1 (Tissue *F* = 1.776, *P* = 0.2013, d.f. = 1; Time *F* = 208.2, *P* < 0.0001 d.f. = 3; Interaction *F* =14.41, *P* < 0.0001, d.f. = 3). GA2ox7 (Tissue *F* = 260.2, *P* = *P* < 0.0001, d.f. = 1; Time *F* = 101.1, *P* < 0.0001 d.f. = 3; Interaction *F* =128.8, *P* < 0.0001, d.f. = 3). A Šídák’s multiple comparisons post hoc test was used for multiple comparisons **p-value < 0.01, ****p-value < 0.0001. d) Length of 3d dark grown hypocotyls after illumination of Col-0 and *ga2oxhept* mutant. Individual data points plotted with a non-linear fit of data 2^nd^ order polynomial and 95% confidence interval r^2^ = 0.1 and 0.3 respectively. Col-0 n=8, *ga2oxhept* n=11. Experiment was completed twice with similar results.

### Limited GA depletion in a *ga2ox* heptuple mutant causes hypocotyl hyperelongation during photomorphogenesis

During the dark to light transition, the distribution of GA in the hypocotyl undergoes reprogramming which we have been able to visualise for the first time using nlsGPS2. We observed a greater decrease in the sub-apical hypocotyl than in the apical hook indicating a potential difference in the response to light in these sections of the hypocotyl. To determine potential mechanisms of these differential regulations, we examined the transcripts of selected metabolic genes in the dark at 1h, 3h and 6h transfer to light in the sub-apical hypocotyl and the apical hook. The selected transcripts examined were those that demonstrated the greatest fold change in the whole hypocotyl at 4 hour transfer to light (Figure S11a). Upon exposure to light, we observed a rapid decline of expression of *GA20ox1* and *GA3ox1* after 1 hour in both the apical hook and the sub-apical hypocotyl followed by an increase limited to the sub-apical hypocotyl at 3 hours (Figure 6c). The latter result could be an indication of a transient or internally localised GA increase in expanding cells during apical hook opening that was missed in our 6 hr analysis of epidermal and cortical cells (Figure 6b). During the dark to light time course, *GA2ox1* expression increased simultaneously in the hypocotyl and the hook until 3 hours (Figure 6c). In contrast, *GA2ox7* expression is strongly increased but this is limited to the apical hook region (Figure 6c).

Previous studies have demonstrated that hypocotyl growth arrest during de-etiolation is dependent on GA levels (Folta et al., 2003). We have shown local and temporally dynamic enzyme expression and GA levels in deetiolating hypocotyls. To establish the importance of reprogramming of GA levels, we observed the growth of the *ga2oxhept* mutant during de-etiolation. The growth rate of Col-0 and *ga2oxhept* mutant diverges at 6 hours after transfer to light with Col-0 growth slowing at a faster rate than *ga2oxhept* (Figure 6d). Taken together, these results indicate that upon transfer to light, reduced biosynthesis and increased catabolism in the hypocotyl leads to decreased GA levels and this depletion is quantitatively important for slowing elongation.

## Discussion

Plant developmental plasticity emerges in part via integration of environmental signals into hormone dynamics, with distinct outcomes based on developmental context. For both embryo protrusion from seed coats in germination and seedling protrusion through soil in skotomorphogenesis, GA accumulation coordinates endogenous signals with environmental cues, particularly light and darkness, to appropriately direct organ growth. In Arabidopsis seed germination, PIF1 antagonizes GA biosynthesis and promotes catabolism (Gabriele et al., 2010; Oh et al., 2004), thus linking light with GA accumulation. Using the direct GA biosensor nlsGPS1, we previously uncovered a GA accumulation gradient in skotomorphogenic hypocotyl cells that was absent in photomorphogenesis (Rizza et al., 2017). It was known that light antagonises GA in this developmental context, but the complexity of light signalling and GA metabolism in Arabidopsis provided multiple potential points of regulation for cellular GA dynamics.

Here we investigated the biochemical basis of this hypocotyl GA gradient and discovered that COP1 signalling acting, presumably indirectly, on the patterned expression of the GA20ox1 biosynthetic enzyme is a central regulatory module. We also show, using a *ga2oxhept* mutant, that GA catabolism contributes to articulating the gradient. Although we previously showed the *pifq* mutant did have lower GA levels in the dark, we now demonstrate that PIFs were dispensable for the GA gradient formation and were instead key for maintenance of skotomorphogenesis. Brassinosteroid signalling has been shown to both promote and repress GA accumulation (Unterholzner et al., 2015; Albertos et al., 2022), but providing epi-brassinolide or BRZ inhibitor did not abolish the GA gradient. As PIF and BR patterning are unlikely to explain GA20ox1 expression in the sub-apical hypocotyl and HY5 activity is limited in the dark, determining the responsible regulators downstream of COP1 will be an interesting topic for a future study.

The discovery of a GA gradient correlated with cell length in etiolated hypocotyls was not evidence of GA levels acting in a simple dose-response relationship with elongation. Rather, overaccumulation of GA in the *ga2oxhept* mutant and after induction of *GA20ox1* expression did not alter cell elongation patterning and thus we can conclude that GA accumulation is not alone sufficient to set cell length in the dark. A similar dynamic was determined in Arabidopsis root elongation in the light (Rizza et al., 2017). Nonetheless, GA is required for subapical hypocotyl cell elongation and a spatially resolved lowering of GA levels will be required to fully reveal the quantitative relationship between GA and the cell elongation gradient. A model in which local GA concentrations cooperate with levels of other growth factors (e.g. PIFs, brassinosteroid) to set cell length would have distinct implications for morphogenesis control than, for example, one in which a low threshold of GA potentiates growth (i.e. via DELLA degradation) with other factors setting cell length. For example, in a cooperative model, a range of GA concentrations would be meaningful and provide robustness for cell length patterning while in a thresholded model GA would provide a gating function for growth.

While relative depletion of GA from the apical hook was not important in skotomorphogenesis, we showed that catabolism of GA was quantitatively important in the early stages of photomorphogenesis. The engineering of GPS2 to have improved orthogonality and reversibility over GPS1 enabled us to better track GA depletion during deetiolation and *ga2oxhept* mutant hypocotyls overelongated after illumination. As either local or temporal GA control is meaningful in many plastic traits, for example axial shoot fate switching in grapevine and strawberry and inflorescence architecture in rice (Wu et al., 2016; Zheng et al., 2018; Tenreira et al., 2017), our study serves as a model for using GPS2 *in vivo* to quantify the relationship between environmentally sensitive cellular GA dynamics and developmental plasticity. Indeed, the nlsGPS2 biosensor was transiently introduced and shown to be functional in several monocot crop species (Dao et al., 2023) and stably introduced into *Medicago truncatula* to clarify where and when roots cells accumulate GA during rhizobium induced nodulation, which had previously been shown to be both promoted and inhibited by GA (Drapek et al., 2023).

## Methods

### Plant material and Growth conditions

WT, mutant and transgenic lines seed in this study were *A. thaliana* ecotype Columbia (Col-0) with the exception of *phyAphyBcry1cry2* in the Landsberg (Ler) background. Seeds were chlorine gas-sterilised and plated on ½ Murashige and Skoog (MS) basal medium (Duchefa Biochemie) with 0.025% MES (Sigma, pH5.7 and 1.2% agar (Duchefa Biochemie). Seeds were stratified in the dark at 4°C for 3 days. Light grown seedlings were placed in a growth chamber with long-day growth conditions (16 h light/8 h dark cycling, temperature cycling 22 °C day/18°C night). For dark grown seedlings plates were transferred to 22°C light for 4 hours before being wrapped in two layers of foil to simulate constant darkness and returned to the growth chamber with long day growth conditions (16h/8h temperature cycling 22°C/18°C).

For β-estradiol induction of pUBQ-XVE-AtGA20ox1, pUBQ-XVE-AtGA20ox1-P2A-AtGA3ox1, and pUBQ-XVE-AtGA3ox1 lines in the dark seeds were plated on a fine mesh (100mu nylon, 38% open area, Normesh) transfer was carried out under green light to ½ MS solid medium supplemented with 2.5µM 17-β-estradiol for the time indicated in the text. Mock induction with DMSO was used as a control. For hormone treated seedlings lines were plated on a fine mesh, 24h after transfer to the growth chamber the mesh was transferred to plates containing the appropriate chemical.

### Generation of Transgenic Arabidopsis Lines

The following nlsGPS1 and nlsGPS2 Arabidopsis lines were generated during this work using the Agrobacterium floral dip method and transformants were selected on ½ MS agar plates containing kanamycin Col-0 nlsGPS2, *cop1-4* nlsGPS1, *ga2oxhept* GPS1 and *phyAphyBcry1cry2* nlsGPS1. To generate *ga2oxhept* homozygous mutant containing null alleles of seven *GA2ox* genes, *ga2ox7-2* (SALK_055721) and *ga2ox8* (SALKseq_040686.2) were crossed to generate *ga2ox7/ga2ox8*. This was then crossed to *ga2oxq*, subsequent generations were backcrossed to *ga2oxq* and until the homozygous line was generated. To generate *cop1-4/hy5-215* nlsGPS1 *cop1-4* nlsGPS1 was crossed to *cop1-4/hy5-215*, the F1 selfed and the homozygous line identified in the F2. Generation of *cop1-4* pUBQ10-XVE-GA20ox1-P2A-GA3ox1 nlsGPS1 was achieved by crossing *cop1-4* to pUBQ10-XVE-GA20ox1-P2A-GA3ox1 nlsGPS1. Homozygous lines were confirmed by using PCR-based genotyping using allele specific primers, those containing nlsGPS1 were screened at seedling stage for fluorescence, those containing inducible lines were screened on hygromycin.

### Engineering and characterization of GPS2 biosensor

Site direct mutagenesis of GPS1 (plasmids pDR-FLIP43 GPS1 and p16-FLIP43 nlsGPS1 (Rizza et al., 2017) was achieved using QuikChange site-directed mutagenesis kit and Phusion high-fidelity DNA polymerase (NEB) following manufacturer protocol resulting in plasmids pDR-FLIP43 GPS2 (for yeast expression) and p16-FLIP43 nlsGPS2 (for plant expression). Expression and fluorescence characterization of biosensors in protease-deficient yeast was performed as described previously (Jones et al., 2014). *Saccharomyces cerevisiae* strain BJ5465 (ATCC 208289 *MATa ura3-52 trp1 leu2-*Δ*1 his3-*Δ*200 pep4:HIS3 prb1-*Δ*1.6 R can1 GAL*) was transformed with plasmid pDR-FLIP43 GPS2 and selected on synthetic complete (SC) medium with 0.8 % agar supplemented with 240 mg/L leucine and 20 mg/L tryptophan (SC agar +Leu, +Trp) for complementation of uracil auxotrophy by the URA3 marker. Transformed yeast was grown in 50 ml SC medium-Ura in 250 ml flasks for two nights. Following, centrifugation, 500 µl – 700 µl chilled silicon bead slurry (MOPs buffer, 0.1% Triton X-100 and 50% (v/v) Zirconia/Silica beads (0.5 mm; Biospec) was added to each tube containing the cell pellets. The tubes were vortexed for 15-20 minutes at 4 °C. The cell lysate was centrifuged at 10000g for 10 minutes at 4 °C. Yeast lysates were diluted 1:2 in 20 mM Tris-HCl, 5 mM imidazole, pH 7.4 and then bound to poly-prep chromatography columns containing His-bind resin (Novagen). The yeast lysate mixture was loaded in the columns, and a 30-minute rolling at 4°C was carried out to allow bonding. Columns were washed twice with 50 mM MOPs, 10 mM imidazole, pH 7.4 and eluted in 50 mM MOPS, 150 mM imidazole, pH 7.4. The elute was diluted with 50 mM MOPS, pH 7.4 in clear-bottom 96-well plates (Greiner) with GA_4_ concentrations as indicated (Sigma) and analysed with a fluorometer (SpectraMax, Molecular Biosciences). Fluorescence readings were acquired for the acceptor FP emission wavelength (Acceptor emission: Am) when samples were excited with the excitation wavelength for acceptor FP (Acceptor excitation: Ax) as the control of sensor expression. An emission scan, including donor and acceptor emission spectra, was acquired when exciting with donor fluorophore excitation wavelength (Donor excitation: Dx). The emission scan range was set between 470 to 550 nm with a step size of 5 nm and excitation at 428 nm. For all imaging, the bandwidth was set to 12 nm, the number of flashes ten, and the integration time 40 µs. BJ5465 yeast containing an empty vector was used as a negative control for background subtraction. GraphPad was used to analyse affinity. Technical replicates were averaged to determine the mean emission ratio for each sample with or without ligand. The intensity of emission of a donor over the intensity of emission of FRET (DxAM/DxDm) was calculated as the emission ratio. Titration curves were analysed using GraphPad software.

### Reversibility determination

Zeba Spin Desalting Columns, 0.5 ml, 40 K MWCO (Thermo Scientific) were used to perform *in vitro* reversibility binding test. Purified sensors were pre-treated with 0 µM GA_4_ and 0.1 µM GA_4_. After two 20-minute buffer exchange steps with 50 mM MOPS pH 7.4 using Zeba spin columns, the eluted GPS2 in 50 mM MOPS pH 7.4 was treated with 0 µM GA_4_ and 0.1 µM GA_4_. Fluorescence emission ratios were analysed as above. in clear-bottom 96-well plate with a SpectraMax fluorometer.

### Phenotypic Characterisation

For hypocotyl length assays plants were grown vertically as described in growth conditions. Plates were imaged after 3d for dark grown hypocotyls and hypocotyls lengths were measured using Fiji software. At least three independent experiments were performed and the sample size is indicated in the figure legends.

### Infrared Imaging and Analysis

For imaging dark grown hypocotyls and de-etiolation, a custom IR imaging setup was used. Images were acquired at 1 hour intervals. At 72h after start of germination light pulse the chamber light was turned on. Hypocotyl lengths and the angles between the cotyledons and the hypocotyl were measured using the Fiji software. At least three independent experiments were performed and the sample size is indicated in the figure legends.

### RNA Extraction and Expression analysis

Seedlings grown for hypocotyl RNA extraction were grown on Normesh in either long day or dark conditions for 3 days. Dark to light transition samples were grown in the dark for three days and transferred to light for four hours before harvest. The hypocotyls were then collected for RNA extraction. Total RNA was extracted using RNeasy Plant Mini Kit (Qiagen). After DNase digest (TURBO™ DNase, Invitrogen) 1µg total RNA was used for cDNA synthesis (SuperScript™ VILO™ cDNA Synthesis Kit, Invitrogen). The expression of genes was tested by qPCR (LightCycler® 480 SYBR Green I, Roche).

Seedlings grown for RNA extraction from the hook and the hypocotyl were grown on a fine strip of Normesh such that the hypocotyl extended above the mesh onto the media. Dissection was carried out following a procedure adapted from a grafting protocol (Melnyk, 2017). Total RNA was extracted using RNAqueous™-Micro Total RNA Isolation Kit (Ambion). After DNase digest (TURBO™ DNase, Invitrogen) 0.5µg total RNA was used for cDNA synthesis (SuperScript™ VILO™ cDNA Synthesis Kit, Invitrogen). The expression of genes was tested by qPCR (LightCycler® 480 SYBR Green I, Roche).

### GUS staining

Etiolated three-day-old seedlings were incubated for 4h at 37°C in the GUS buffer (200mM NaH2PO4, pH7.0, 0.5M EDTA pH8.0, 0.1% Triton-X, 20mM Potassium ferricyanide, and 20mM Potassium ferrocynanide containing 20mM 5-bromo-4-cholro-3-indolyl glucuronide. To stop the GUS reaction, the reaction mix was replaced with an ethanol concentration series (30 min minimum) at 30%, 50%, 70%, and 100%. Seedlings were imaged with a Keyence microscope.

### Confocal imaging

Seedlings were mounted in liquid ¼ x MS medium (1/4 X MS salts, 0.025% MES, pH5.7), covered with a coverslip and imaged. Confocal images were acquired with a format of 1024×524 pixels and a resolution of 12 bit on an upright Leica SP8FLIMan using a 10x dry objective for dark grown hypocotyls and a 20x dry 0.70 HC PLAN APO objective for light grown hypocotyls. To excite Cerulean and Aphrodite, 448 nm and 514 nm lasers were used, respectively. The 552 nm laser line was used to excite propidium iodide (PI, Sigma-Aldrich). Emission filters were 460 to 500 nm for Cerulean, 525 to 560 nm for Aphrodite, and 590 to 635 nm for PI. Three fluorescence channels were collected for FRET imaging: Cerulean donor excitation and emission or DxDm, Cerulean donor excitation, Aphrodite acceptor emission or DxAm, and Aphrodite acceptor excitation and emission or AxAm. The laser power was set to 3% to excite Cerulean and 2% to excite Aphrodite with detector gain set to 110.

### Imaging Processing and Analysis

Imaging process and analysis were performed with FRETENATOR plugins (Rowe et al., 2022) Segmentation settings were optimised for each experiment but kept constant within each experiment. The AxAm channel was used for segmentation. For segmentation Otsu thresholds were used, difference of Gaussian kernel size was determined empirically and a minimum ROI size was set to 20. Distance from meristem was defined using FRETANATOR ROI labeller.

### Accession Numbers

N2110211: *ga20ox1,ga20ox2,ga20ox3* nlsGPS1; N2110213: pUBQ10-XVE-GA20ox1 nlsGPS1; N2110214: pUBQ10-XVE-GA3ox1 nlsGPS1; N2110216: pUBQ10-XVE-GA20ox1-P2A-GA3ox1 nlsGPS1; N2107737: *pifq* nlsGPS1; N2107734: Col-0 nlsGPS1

## Acknowledgements

We thank P. Hedden, S.G. Thomas, A. Phillips, R. Ulm, J.J. Casal and T.P. Sun for kindly providing us *cop1-4*, *cop1-4/hy5-215, ga2oxq* (Rieu et al., 2008), *ga20ox1/2/3,* pGA20ox-GUS (Plackett et al., 2012), pGA3ox-GUS (Mitchum et al., 2006) and *phyAphyBcry1cry2* (Mazzella et al., 2001). Martin Balcerowicz for comments on the paper. R. Wightman, G. Evans and L. Tully for help with microscopy and horticulture. This work was supported by the Gatsby Charitable trust (GAT3395) to AMJ and European Research Council under the European Union’s Horizon 2020 research and innovation program (grant agreement n° 759282) to AMJ.

## Author Contributions

J.G., A.R., B.T. and A.M.J. performed experiments. J.G., A.R., B.T. and A.M.J. analysed the results. J.G. and A.M.J. wrote the manuscript. J.G, B.T, A.R., and A.M.J. revised the manuscript. W.B.F. and A.M.J. secured funding. A.M.J. and W.B.F. conceived the engineering of GPS2. J.G. and A.M.J. designed the research. A.M.J. supervised the project.

---

**Figure S1.**
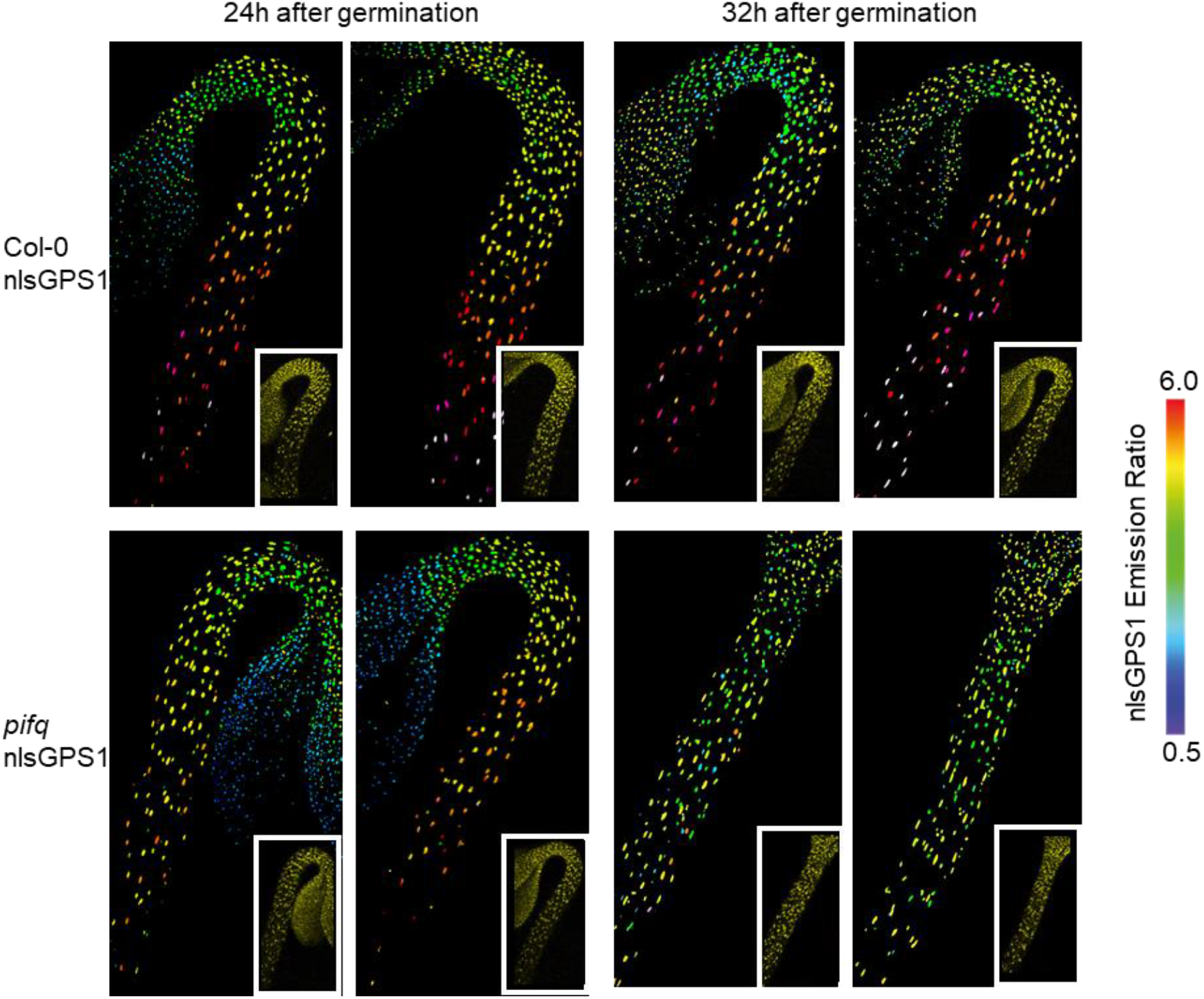
Additional images of Col-0 nlsGPS1 and *pifq* nlsGPS1 dark grown hypocotyls. nlsGPS1 emission ratios of dark grown hypocotyls 24h and 32h after germination. Representative images of emission ratios and YFP fluorescence (Inset) are shown (Images corresponding to emission ratio images presented in Figure 1d).

**Figure S2.**
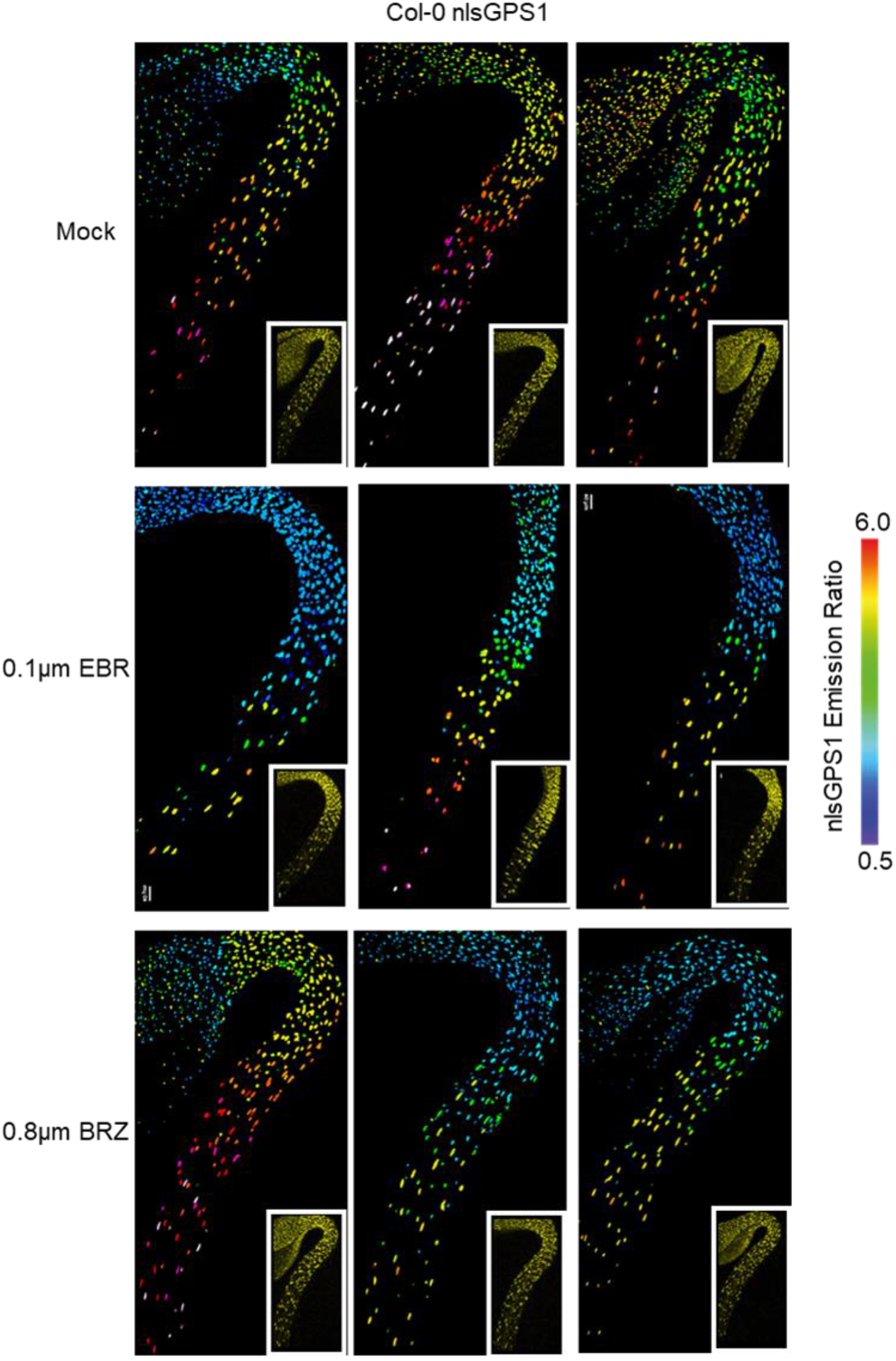
Additional images of Col-0 nlsGPS1 dark grown hypocotyls treated with EBR and BRZ. Emission ratios and YFP images (inset) of dark grown hypocotyls treated with 0.1µm epi-brasssonolide or 0.8µm brassinazole for 24h before imaging with DMSO treatment as mock. (Images corresponding to emission ratio images presented in Figure 1g)

**Figure S3.**
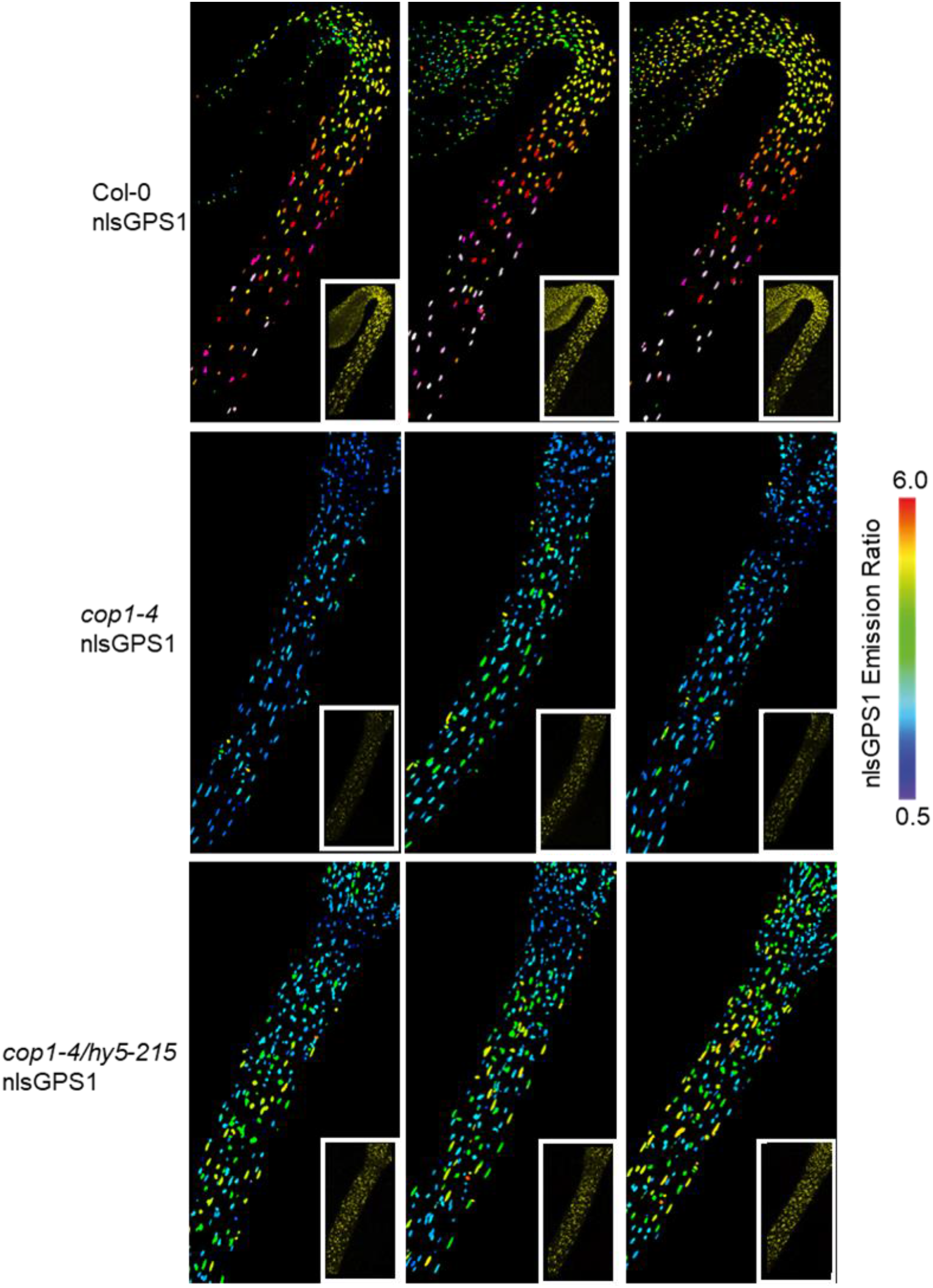
Additional images of Col-0 nlsGPS1, *cop1-4* nlsGPS1 and *cop1-4/hy5-215* nlsGPS1 dark grown hypocotyls. Emission ratios and YFP images (inset) of 3d dark grown hypocotyls. (Images corresponding to emission ratio images presented in Figure 2d.)

**Figure S4.**
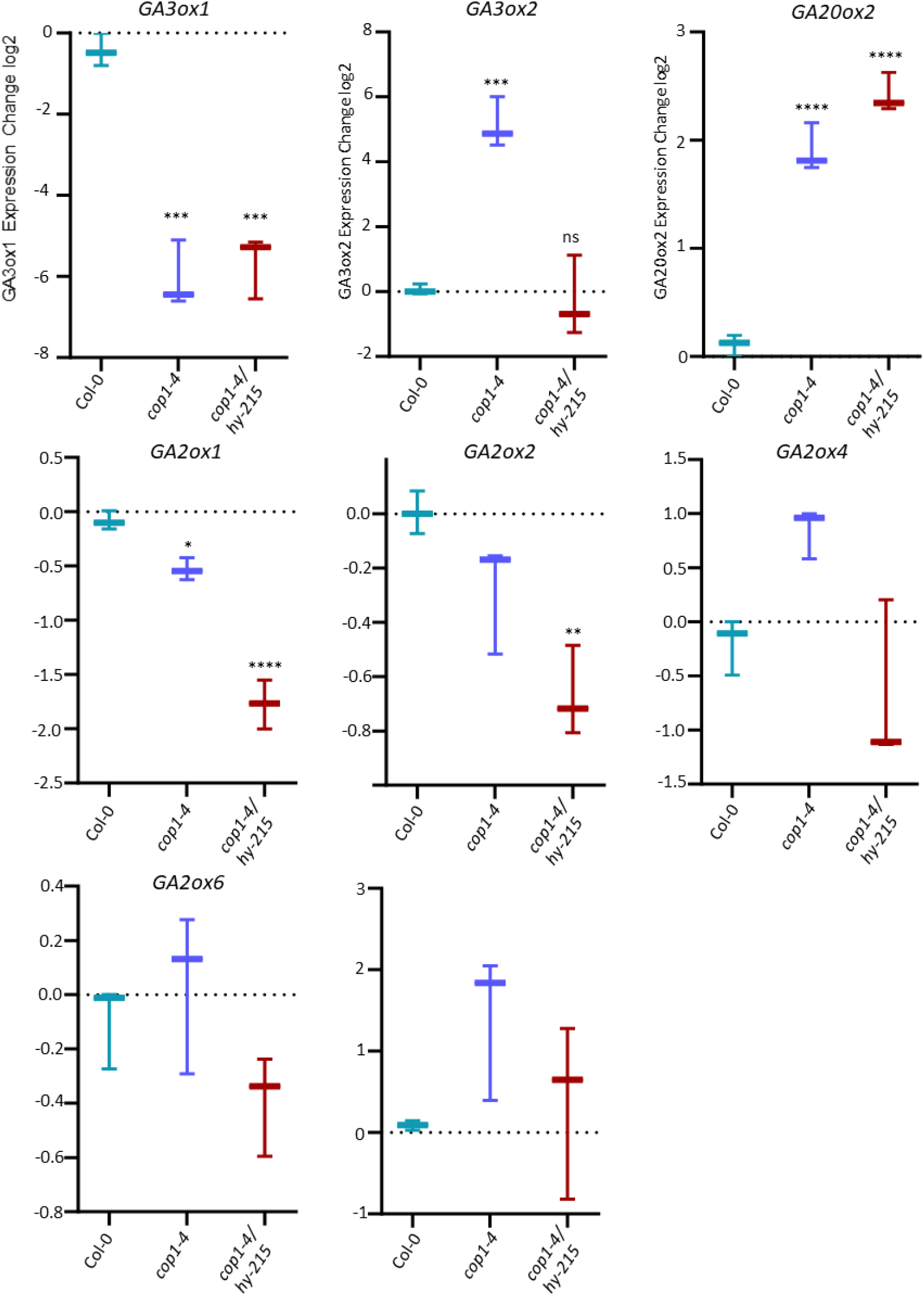
Relative expression of GA metabolic genes in Col-0, *cop1-4* and *cop1-4/hy5-215* dark grown hypocotyls. Log2 fold change from Col-0 for *cop1-4* and *cop1-4/hy5-215* are shown. Reference gene P2AA3 (At1g13320). A one-way ANOVA was performed to compare the effect of genotype on gene expression. A Dunnet’s post hoc test was used for multiple comparisons to Col-0 control * p-value <0.05, **p-value < 0.01, ***p-value < 0.001, ****p-value < 0.0001

**Figure S5.**
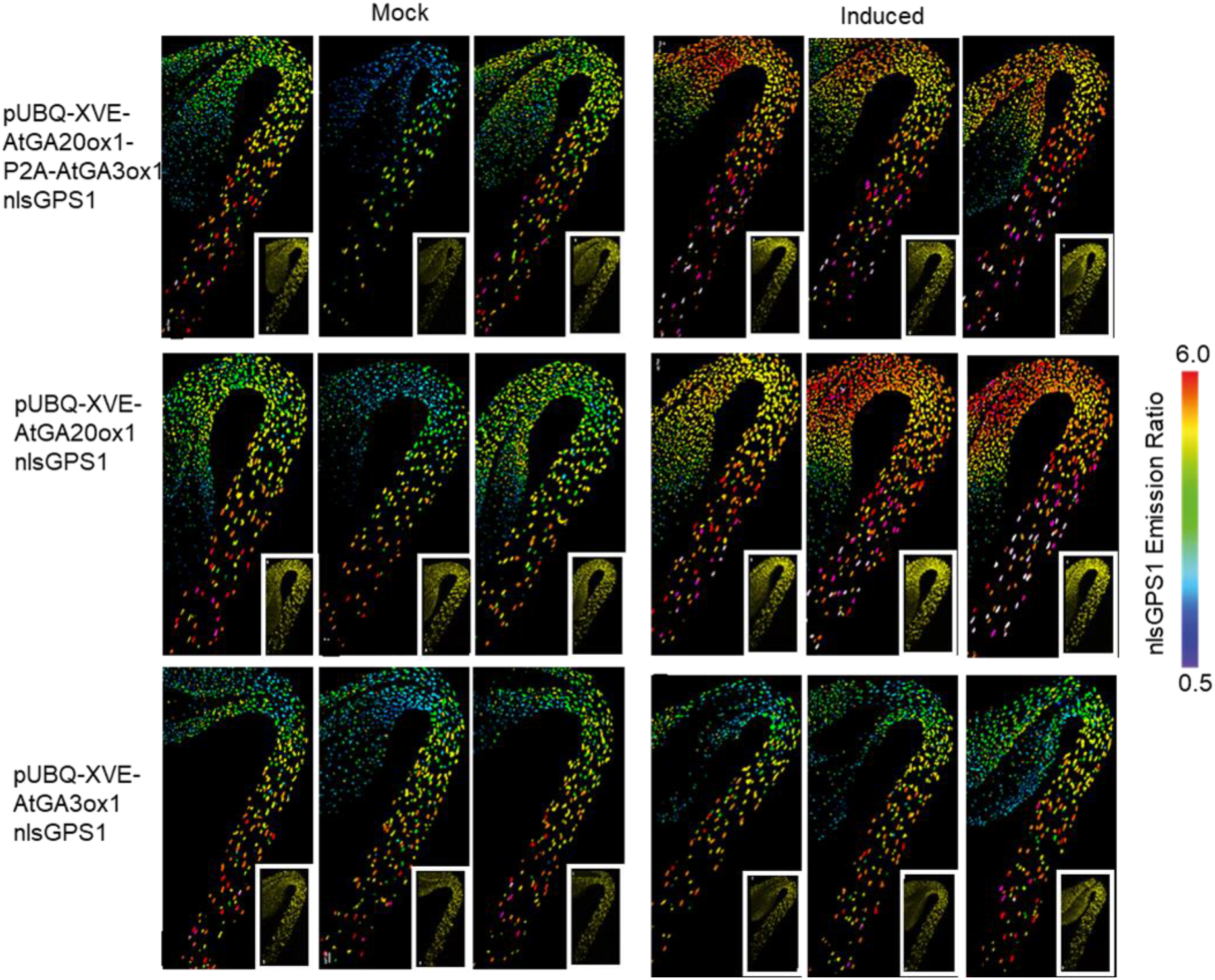
Additional images of GA enzyme induction in Col-0 nlsGPS1. Emission ratios and YFP images (inset) of dark grown hypocotyls. β-estradiol inducible GA enzyme transgenic lines 48 h after induction with 2.5 µM 17-β-estradiol (induced) or with 0.1% DMSO mock induction (mock). (Images corresponding to emission ratio images presented in Figures 3g-i)

**Figure S6.**
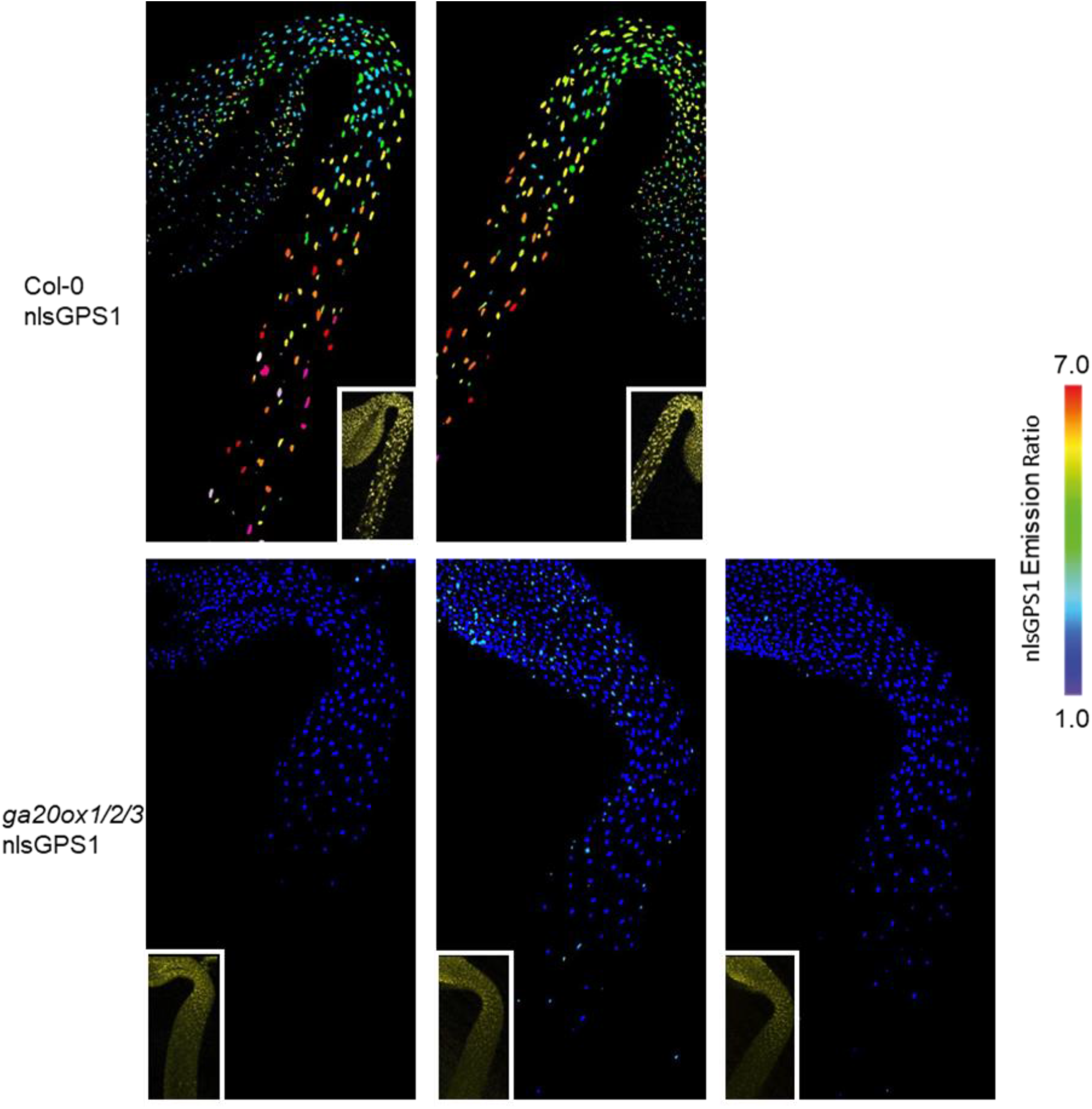
Additional images of Col-0 nlsGPS1 and *ga20ox1/2/3* nlsGPS1 dark grown hypocotyls. Emission ratios and YFP images (inset) of 3d dark grown hypocotyls. (Images corresponding to emission ratio images presented in Figure 4c.)

**Figure S7.**
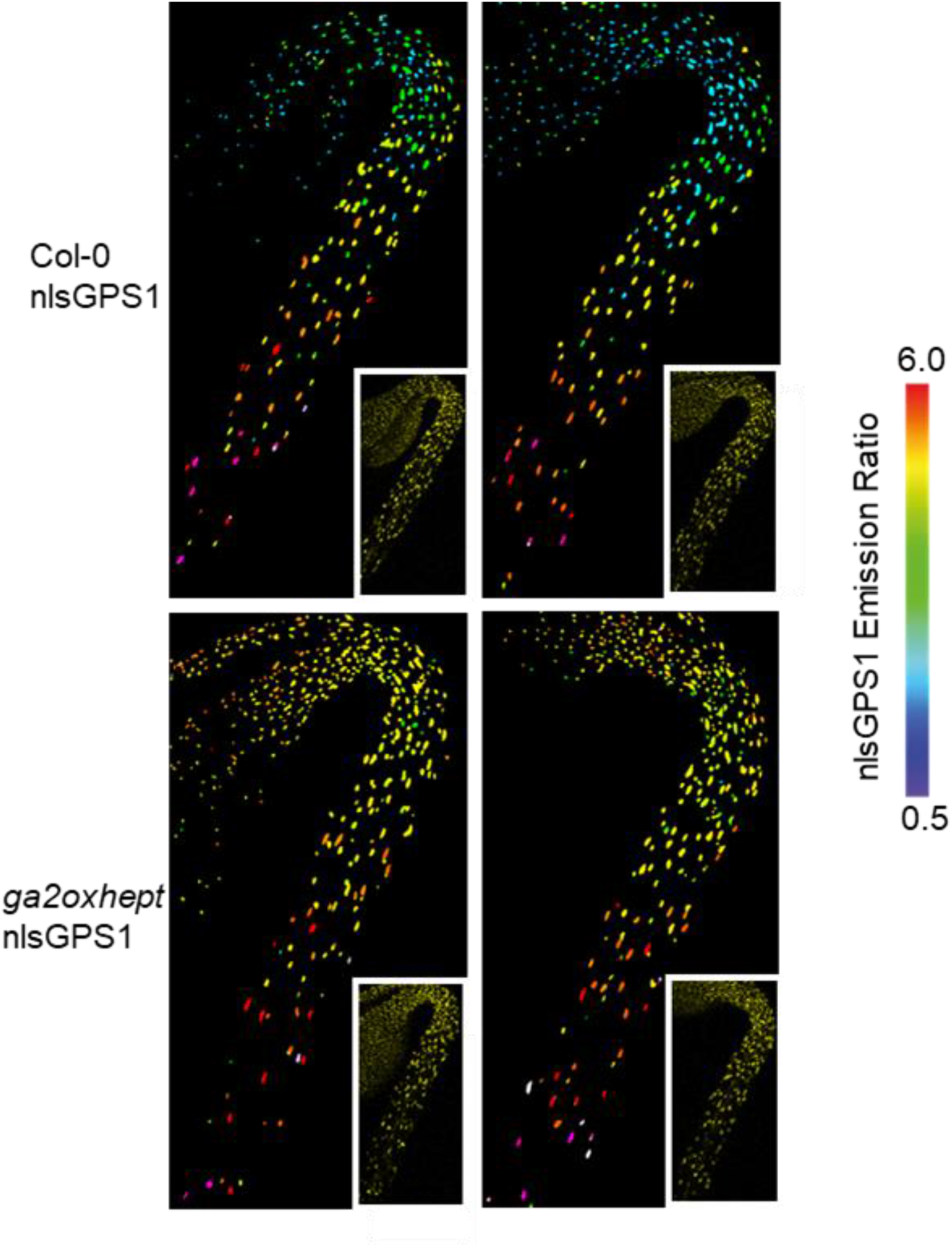
Additional images of Col-0 nlsGPS1 and *ga2oxhept* nlsGPS1 dark grown hypocotyls. Emission ratios and YFP images (inset) of 3d dark grown hypocotyls. (Images corresponding to emission ratio images presented in Figure 4e.)

**Figure S8.**
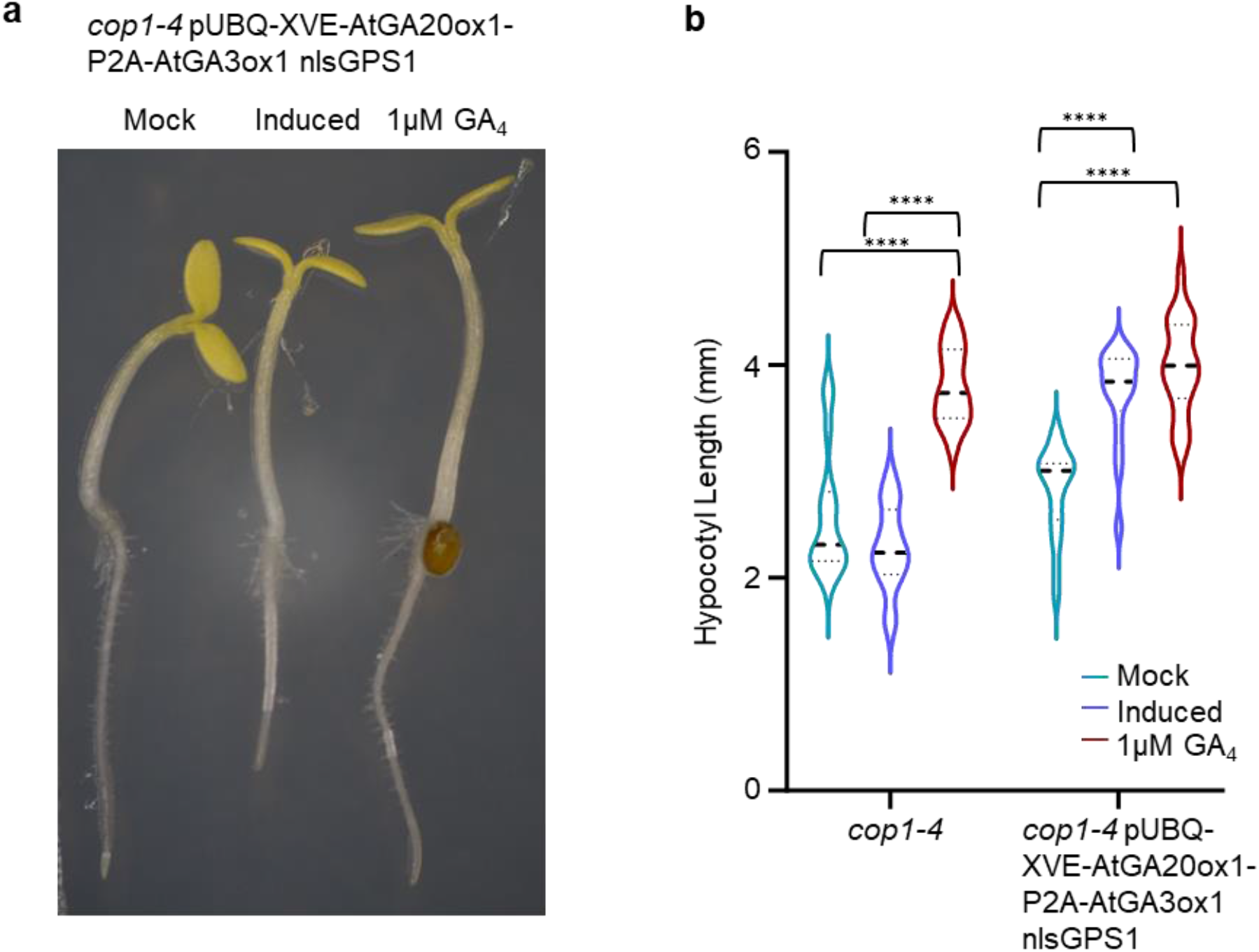
Hypocotyl length after GA enzyme induction in *cop1-4* nlsGPS1. a) Representative images of *cop1-4* pUBQ-XVE-AtGA20ox1-P2A-AtGA3ox1 nlsGPS1 3 day old dark grown hypocotyls 48 h after mock induction with 0.1% DMSO (mock), induction with 2.5 µM 17-β-estradiol or on media containing 1µM GA_4_. b) Hypocotyl lengths of *cop1-4* and *cop1-4* pUBQ-XVE-AtGA20ox1-P2A-AtGA3ox1 nlsGPS1 3d dark grown hypocotyls 48 h after mock induction with 0.1% DMSO (mock), induction with 2.5 µM 17-β-estradiol or on media containing 1µM GA_4_. Two-way ANOVA (Treatment F = 68.55, P <0.0001 d.f. = 2; Genotype F = 51.24, P < 0.0001 d.f. = 1; Interaction F = 21.12, P = < 0.0001, d.f. = 2. (n> 16 biologically independent hypocotyls for all genotypes and treatments). A Tukey’s multiple comparisons post hoc test was used for multiple comparisons ****p-value < 0.0001.

**Figure S9.**
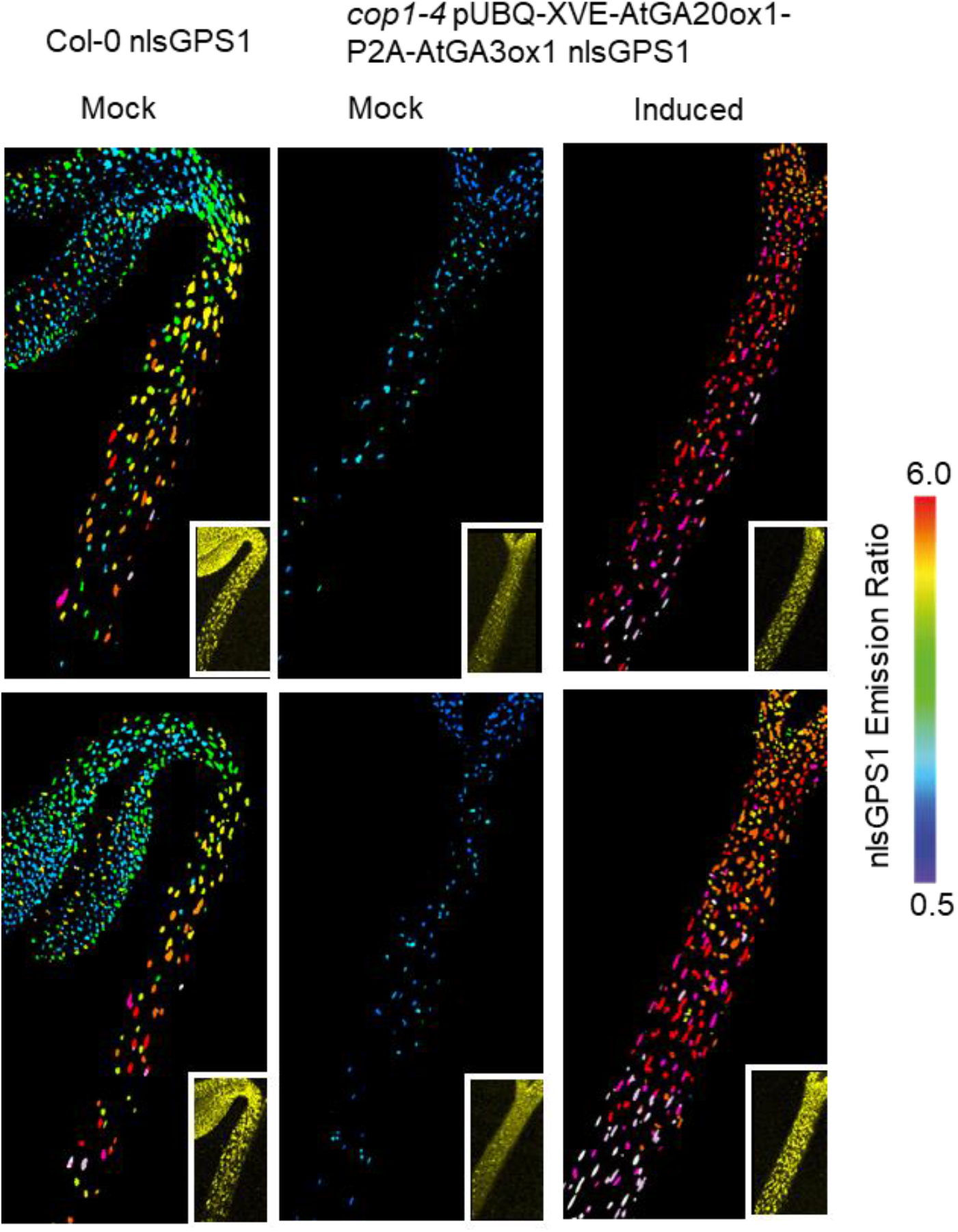
Additional images of GA enzyme induction in *cop1-4* nlsGPS1. Emission ratios and YFP images (inset) of 3d dark grown hypocotyls. Images corresponding to emission ratio images presented in Figure 4H. β-estradiol inducible GA enzyme transgenic lines 48 h after induction with 2.5 µM 17-β-estradiol (induced) or with 0.1% DMSO mock induction (mock). (Images corresponding to emission ratio images presented in Figure 4h)

**Figure S10.**
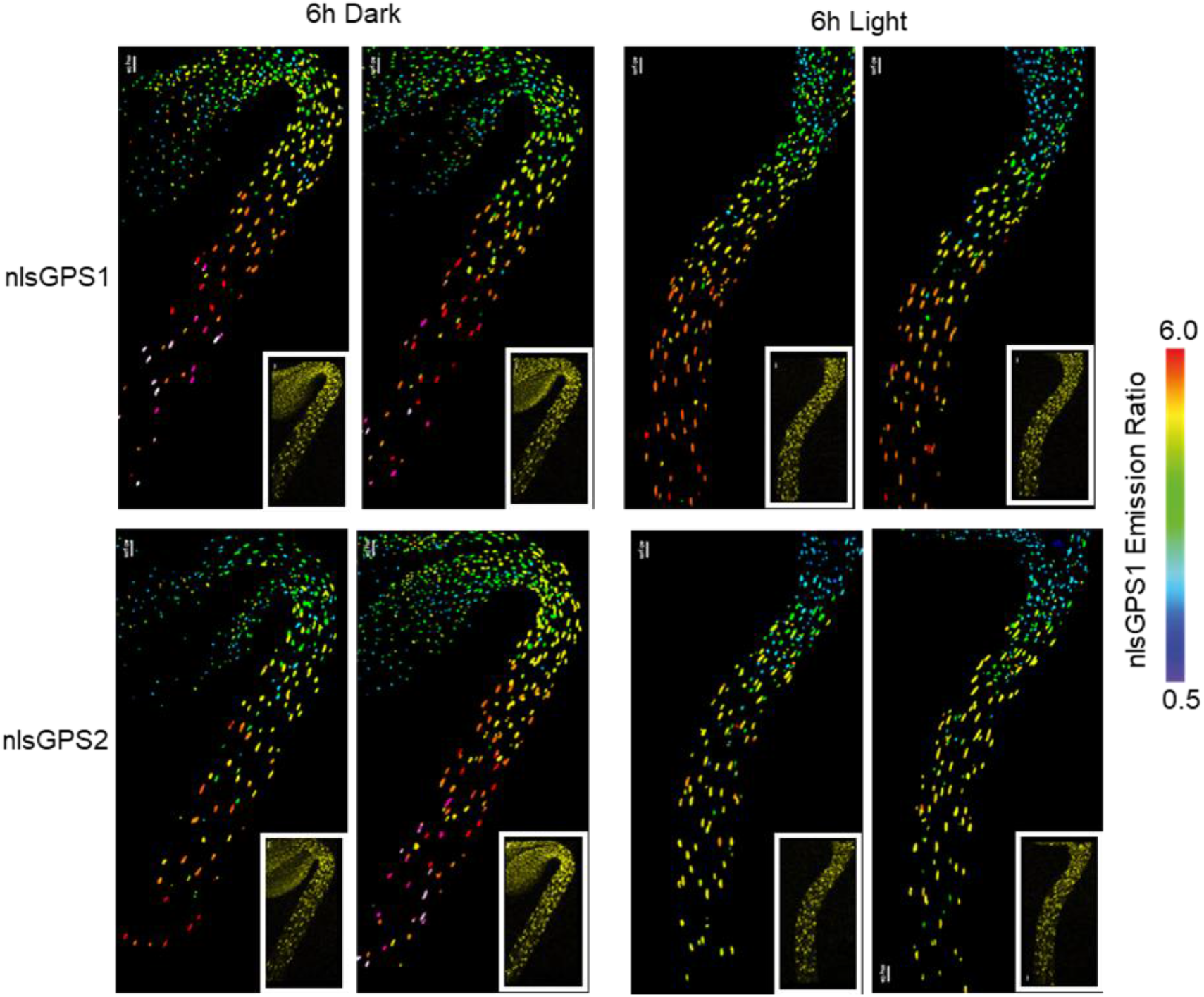
Additional images of dark to light transition nlsGPS1 vs nlsGPS2. Emission ratios and YFP images (inset) of 3d dark grown hypocotyls. (Images corresponding to emission ratio images presented in Figure 6b.)

**Figure S11.**
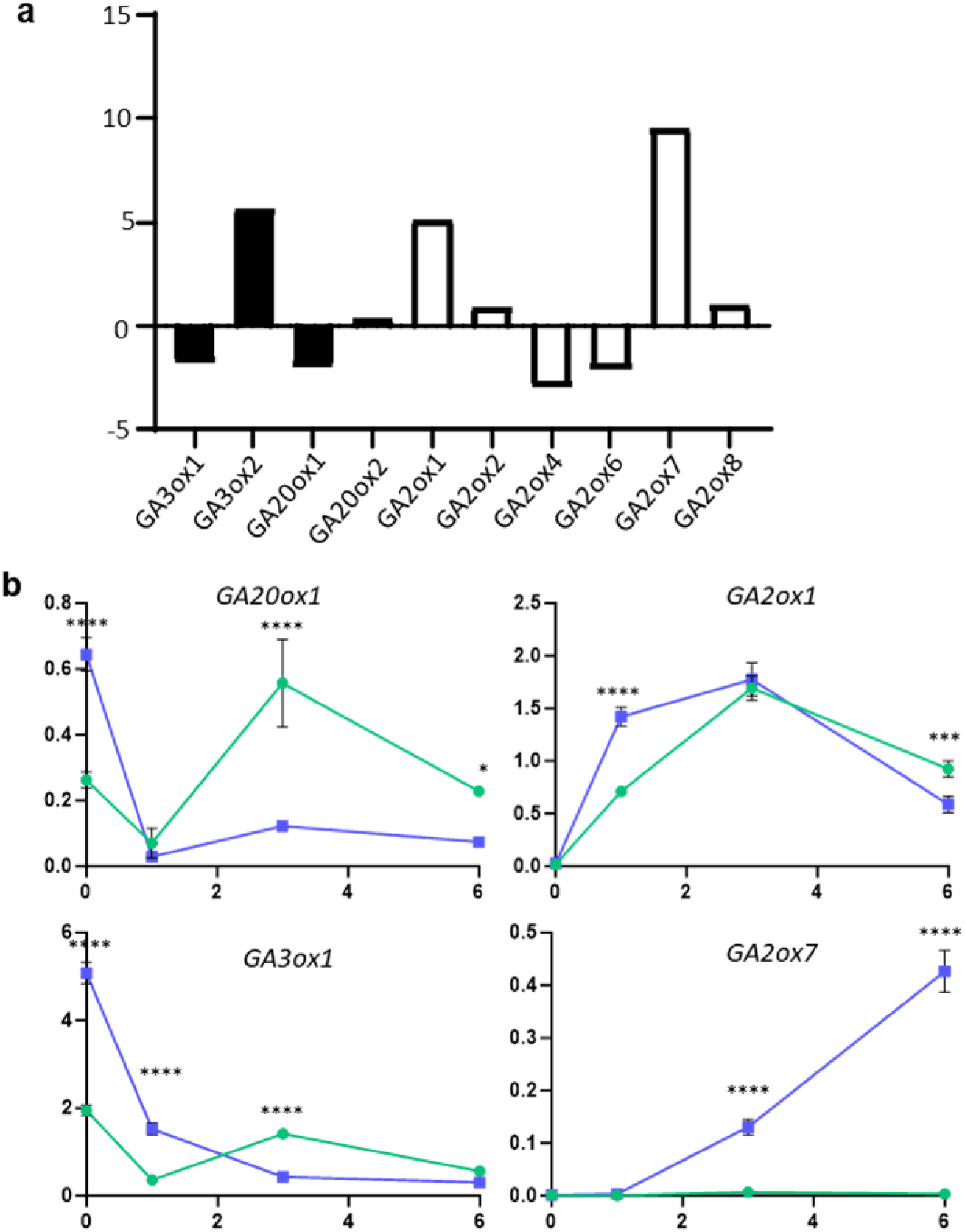
Gene expression in hypocotyls after transfer to light. a) Gene expression change in 3d dark grown hypocotyls transferred to the light for four hours b) Independent replicate of relative expression changes of selected GA metabolic enzymes in the hook and sub-apical hook of Col-0 during de-etiolation shown in figure 6c. Dissections were carried out in the dark and at 1h, 3h and 6h light exposure. Data represents mean and standard deviation of three technical repeats from one biological repeat containing (30) dissected hypocotyls per sample. Two-way ANOVA: *GA20ox1* (Tissue *F* = 7.974, *P* = 0.0122 d.f. = 1; Time *F* = 68.42, *P* < 0.0001 d.f. = 3; Interaction *F* =59.30, *P* < 0.0001, d.f. = 3). *GA3ox1* (Tissue *F* = 266.7, *P* < 0.0001, d.f. = 1; Time *F* = 887.3, *P* < 0.0001 d.f. = 3; Interaction *F* =375.1, *P* < 0.0001, d.f. = 3). *GA2ox1* (Tissue *F* = 10.79, *P* = 0.0047, d.f. = 1; Time *F* = 404.8, *P* < 0.0001 d.f. = 3; Interaction *F* =38.10, *P* < 0.0001, d.f. = 3). *GA2ox7* (Tissue *F* = 391.4, *P* = *P* < 0.0001, d.f. = 1; Time *F* = 218.8, *P* < 0.0001 d.f. = 3; Interaction *F* =214.4, *P* < 0.0001, d.f. = 3). A Šídák’s multiple comparisons post hoc test was used for multiple comparisons *p-value < 0.05, ***p-value < 0.001, ****p-value < 0.0001

